# Transcription regulates the spatio-temporal dynamics of genes through micro-compartmentalization

**DOI:** 10.1101/2023.07.18.549489

**Authors:** Hossein Salari, Geneviève Fourel, Daniel Jost

## Abstract

Although our understanding of the involvement of heterochromatin architectural factors in shaping nuclear organization is improving, there is still ongoing debate regarding the role of active genes in this process. In this study, we utilize publicly-available Micro-C data from mouse embryonic stem cells to investigate the relationship between gene transcription and 3D gene folding. Our analysis uncovers a nonmonotonic - globally positive - correlation between intragenic contact density and Pol II occupancy, independent of cohesin-based loop extrusion. Through the development of a biophysical model integrating the role of transcription dynamics within a polymer model of chromosome organization, we demonstrate that Pol II-mediated attractive interactions with limited valency between transcribed regions yield quantitative predictions consistent with chromosome-conformation-capture and live-imaging experiments. Our work provides compelling evidence that transcriptional activity shapes the 4D genome through Pol II-mediated micro-compartmentalization.

## Introduction

Chromosome conformation assays like Hi-C unveiled hierarchical organization of chromosomes within eukaryotic nuclei ^1,2^. In metazoans, Mbp-scale “checkerboard” patterns in contact maps reveal spatial segregation of chromosomes into a euchromatic “A” compartment and a heterochromatic “B” compartment ^3,4^. At a smaller scale (∼100s kbp), chromosomes fold into topologically associating domains (TADs) and loops ^5–7^. The prevailing model ^8^ suggests that compartments emerge from the micro-phase separation of epigenomic regions mediated by chromatin-binding architectural proteins ^9,10^, while most TADs result from cohesin-driven chromatin loop extrusion with CTCF acting as a barrier ^11,12^. Hi-C and live imaging experiments indicate that depleting CTCF or cohesin disrupts TADs, weakens CTCF-mediated loops, but has limited effects on compartmentalization ^7,13–16^. Nevertheless, some loops or TADs remain unaffected by these treatments ^14,17^, likely originating from distinct mechanisms.

3D chromosome organization regulates gene expression during interphase ^18–20^. Notably, colocalization of promoters and enhancers within TADs can directly influence transcription initiation, potentially increasing transcription rates ^20,21^. Conversely, recent studies suggest that genes serve as central units of the 3D genome and that transcription itself plays a role ^22–25^. High-resolution contact maps in mammalian and fly cells reveal transcription-dependent fine structures, such as loops between active gene promoters, promoters and enhancers, or transcriptional start (TSS) and termination (TTS) sites of the same gene ^13,26–29^. However, the mechanistic origins of these fine structures, despite their potential significance in gene regulation, remain controversial.

Indeed, on the one hand, some experiments in Drosophila and mice indicate higher 3D contacts within expressed gene bodies compared to repressed ones ^29,30^. Remodeling of chromatin structure around genes during mouse thymocyte maturation often coincides with transcriptional changes ^31^. RNA Polymerase IIs (Pol II) form also distinct foci and higher-order clusters known as transcription factories ^32–35^, and active genes tend to colocalize within transcriptionally-active subcompartments ^27,36^. On the other hand, there are cases in mammalian cells where significant unfolding of genes occurs after strong transcriptional activation ^37–39^, and acute depletion of Pol I, II, and III has minimal effects on large-scale genome folding ^40^. In budding yeast, gene activity inversely correlates with local chromatin compaction ^25^. Live-cell imaging experiments highlight the relationship between gene transcription and chromatin dynamics ^41–45^, revealing enhanced gene mobility upon Pol II elongation inhibition ^41,43,44^ or gene activation ^42^ and correlated motions between active regions ^44,45^.

Complementing experiments, biophysical and polymer models have also explored the complex interplay between transcription and genome dynamics ^35,37,46–49^. Elongating or backtracked Pol IIs may act as barriers for SMC-mediated extrusion, indirectly impacting genome organization ^38,46,47,50–54^. Transcription-dependent changes in the local chromatin fiber rigidity and contour length may lead to extended conformations for highly-transcribed genes ^37^. Computational models considering Pol II-mediated interactions ^35^, interactions with transcription factories or condensates ^48,49^, or P-TEFb interactions ^43^ suggest that attractive interactions between active regions may capture inter-gene contacts observed in Hi-C and the gene mobility observed in live-imaging.

Overall, the evidence presents a complex understanding of the genome spatio-temporal dynamics in response to gene transcription, necessitating a comprehensive framework to reconcile these observations. In this study, we analyze publicly-available Micro-C data for mouse embryonic stem cells (mESCs) and develop observables to characterize transcription-dependent 3D gene folding. Our analysis reveals a nonmonotonic relationship between intragenic contact density and gene transcription, potentially reconciling contradictory data. By dissecting the contributions of loop extrusion and transcription-associated factors, we propose Pol II occupancy as a key determinant of gene folding. Using a traffic model for gene activity and a 3D polymer model ^55,56^, we demonstrate that transcriptionally-active subcompartments and intragenic contact enrichment may arise from Pol II-mediated phase separation. Furthermore, we suggest that Pol II-mediated condensation, coupled with transcriptional bursting, may slow down gene mobility, aligning with experimental observations.

## Results

### RNA Pol II occupancy and gene length correlate with intra-gene condensation

In this study, we aimed to investigate the potential role of transcriptional activity in the local organization of genes within the genome. To accomplish this, we focused on mouse embryonic stem cells (mESC) as they provide abundant quantitative data. Specifically, we utilized publicly available high-resolution Micro-C and Pol II ChIP-seq data ^13^. For our analysis, Micro-C contact maps were distance-normalized to examine contact enrichment compared to a sequence-averaged null behavior, resulting in the observed over expected (obs/exp) contact map. We introduced two scores for each gene (**Fig. 1A**, **Materials and Methods**): (i) Intra-gene contact enrichment (IC), which represents the mean obs/exp values calculated for all pairs of loci within the gene, capturing the level of self-association and overall gene condensation. (ii) Intra-gene RNA Pol II enrichment (IR), which corresponds to the mean normalized Pol II ChIP-seq profile within the gene and reflects gene transcriptional activity, correlating with RNA-seq data (**Fig.S1**).

**Fig. 1.**
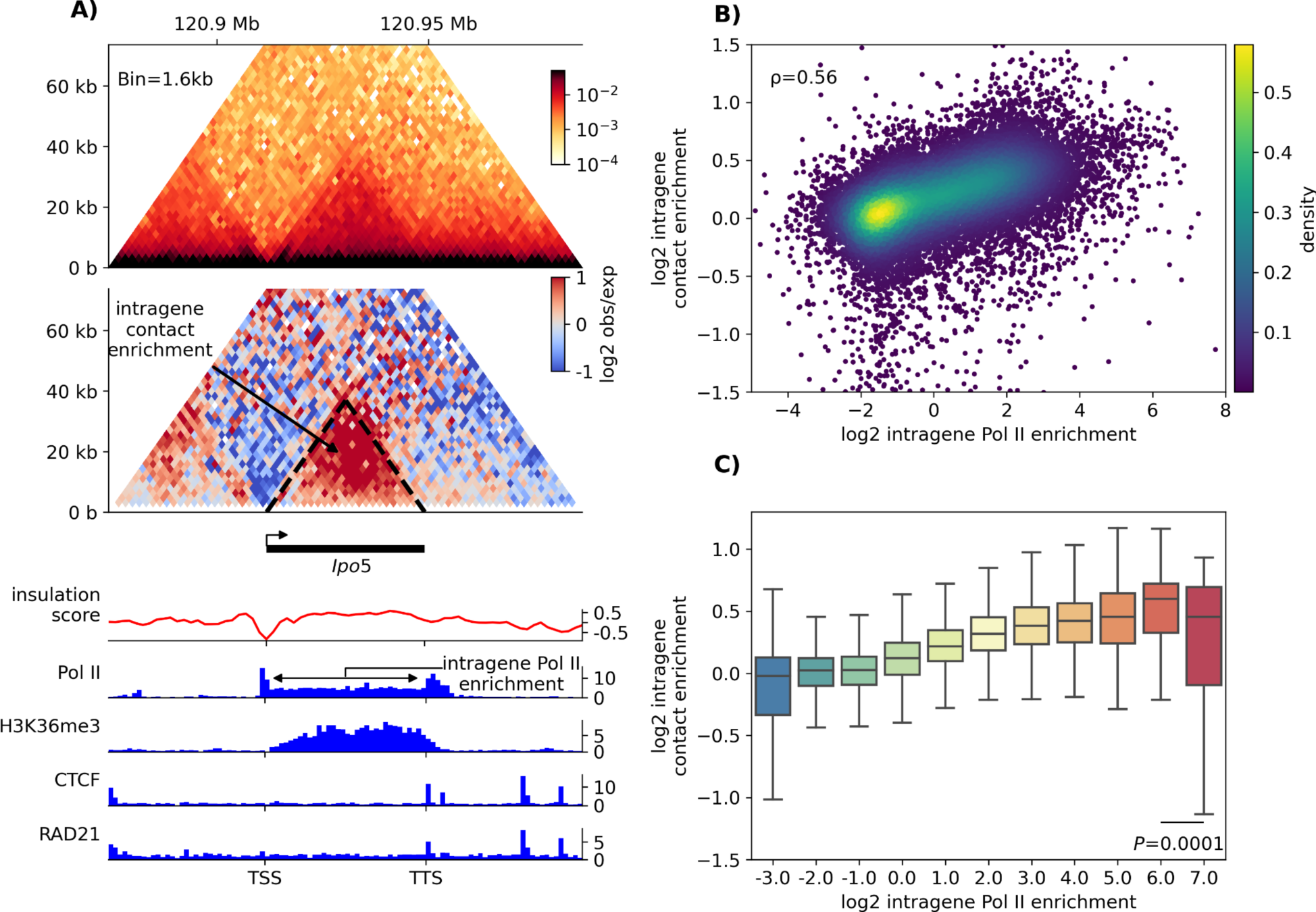
Correlation between intra-gene contact and intra-gene Pol II enrichments. (A) Observed Micro-C contact map (top), observed/expected map (middle) and several ChIP-seq profiles (bottom) of the genomic region including the *Ipo5* gene (chr14:120,874-120,984kb) in mESC. The intra-gene contact (IC) and RNA Pol II (IR) enrichments are illustrated. (B) Scatterplot of IC versus IR for all genes longer than 1kb (24,363 genes). Colors refer to the density of dots. (C) Boxplots of IC after clustering together the genes (dots in (B)) with similar IR.

**Figure 1B** shows a significant positive correlation (Spearman’s ρ = 0.56, t-test *p*-value <1e-200) between IC and IR scores, indicating that increased transcriptional activity is associated with enhanced intra-gene condensation. This result is consistent with prior research on mouse ESC ^29^, *Drosophila* ^30^ and mouse DP and DN3 thymocytes ^31^. We also checked that such a correlation remains mainly independent of the phosphorylation status (Ser5P and Ser2P) of Pol II (**Fig.S2**), which is associated with different dynamical and interacting states of the polymerase ^57^. Additionally, a similar correlation between IC and intra-gene H3K36me3 (a histone mark related to Pol II elongation) content was detected (**Fig.S1**). As a control, we observed weak, negative correlations between IC and repressive marks (H3K27me3 and H3K9me3) (**Fig.S3**). Clustering genes based on similar IR scores revealed a nonlinear and non-monotonic relationship (Fig. 1C): IC generally increases with IR, except at very high Pol II levels where a slight but significant relative decrease in contact frequency occurs. Importantly, this behavior cannot be solely attributed to the inherent properties of Micro-C experiments to detect more or less contacts depending on the molecular crowding on DNA ^25^ since similar behavior was observed using mESC Hi-C data ^19^ (**Fig.S4**). Interestingly, this correlation between IR and IC holds true regardless of gene compartment (A or B) (**Fig.S5**) or the number of exons ^19^ (**Fig.S6**). However, genes with higher exon counts tend to exhibit more intra-gene contacts compared to those with fewer exons. Moreover, IC scores for A-compartment genes are generally higher than those for B-compartment genes, which may suggest an interference between the segregation of heterochromatin and the condensation of Pol II-enriched genes. Interestingly, this difference becomes more pronounced for genes with high IR scores, where the drop in IC is more significant for B-genes.

In mammals, highly active genes are typically smaller, and larger genes, when active, are usually lowly expressed (**Fig.S7**). Hence, we investigated whether gene size may be a confounding factor or, on the contrary, could be a determining factor by classifying genes based on both their IR and genomic length (**Materials and Methods**). Figure 2B displays the average IC score for each category, revealing a positive correlation between IC and gene length: longer genes exhibit stronger intra-gene contact frequency at a given Pol II occupancy density. These findings suggest a cooperative effect in intra-gene folding, wherein both Pol II density and gene size play integral roles ^58^.

**Fig. 2.**
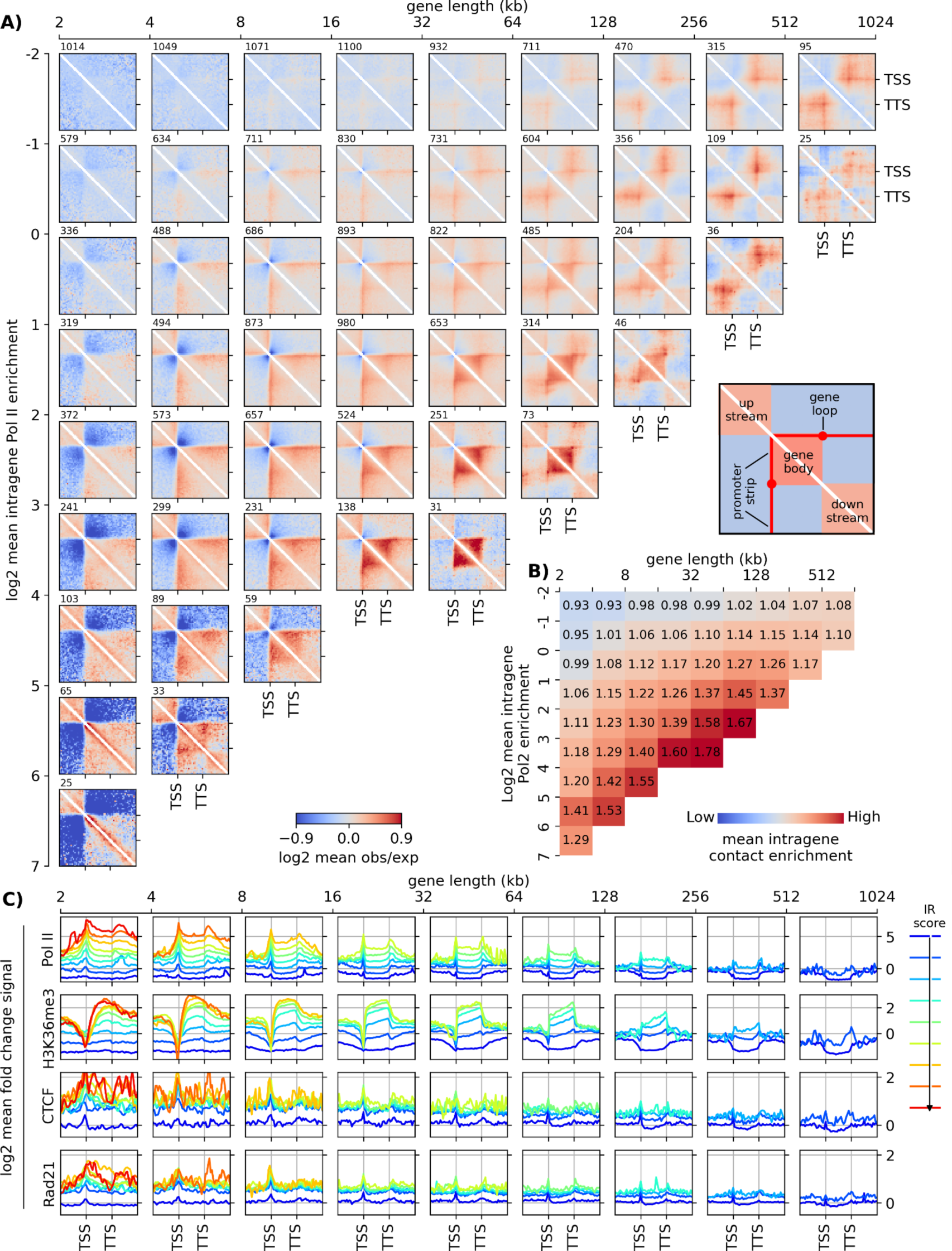
Gene classification based on size and Pol II occupancy, and pileup meta-gene analysis. (A) Pileup meta-gene analysis (PMGA, see **Methods**) of the obs/exp map around genes clustered based on their length (horizontal axis) and Pol II enrichment (vertical axis). The number of genes of each cluster is indicated above on each map. Maps for clusters with less than 25 representative genes were not drawn, due to lack of statistics. (B) Average IC scores for each cluster in (A). (C) PMGA of different chromatin tracks: in each subplot, all the average profiles of the different Pol II clusters for genes of the same length range are shown (from left to right: from small to large genes); different colors correspond to the different Pol II clusters, from low (blue) to high (red) IR score.

To gain deeper insights into the contact patterns and profiles within and surrounding genes, we conducted a pile-up meta-gene analysis (PMGA), aggregating the rescaled obs/exp maps and ChIP-seq profiles of genes with similar size and transcriptional activity (Fig. 2A**,C**, **Materials and Methods**). PMGA uncovered a strong correlation between Pol II profiles and certain structural features of contact maps: intra-gene contact maps were nearly uniform, consistent with the constant Pol II levels observed within genes; stripes of preferential interactions were observed between Pol II-rich promoters/TSSs and gene bodies (“stripe”); and loops were formed between Pol II-rich TSSs and TTSs (“TSS-TTS loops”). Notably, the correlation between Ser2P Poll II (which have no peak at TSS), Ser5P Pol II (only weak peaks at TSS and TTS) and H3K36me3 (strong depletion around TSSs) profiles, taken individually, and Micro-C patterns such as TSS-TTS loops and stripes was less apparent (Fig. 2A**,C****, Fig. S2**). Regarding the dependency on gene size, we found that promoters of short genes are often located at the domain borders, while larger genes tend to form their own insulated domains separate from surrounding regions.

To further investigate the role of Pol II occupancy, we analyzed two publicly available datasets involving the treatment of mESC cells with transcriptional inhibitor drugs: triptolide (TRP), which inhibits Pol II initiation, and flavopiridol (FLV), which inhibits Pol II elongation ^29^. Firstly, we confirmed the significant reduction in the intensity of Pol II-mediated loops after both treatments (**Fig.S8**). Consistent with the observed loss of intra-gene Pol II occupancy in all genes, particularly highly transcribed ones (**Fig.S9,S10**), intra-gene interactions were weaker in the TRP and FLV cases compared to the normal condition (Fig. 3A,**S11,S12**), resulting in a 12% reduction in IC for large active genes post-treatment. These results align with a previously reported observation of 25% reduction in the intensities of gene stripes following Pol II inhibition ^29^. Moreover, there exists a notable correlation between the fold-changes (treated vs untreated) in IC and IR scores (Fig. 3C): the greater the reduction in Pol II occupancy for a given gene, the more likely its intra-gene folding is affected. Interestingly, when re-clustering genes based on their new IR scores measured in TRP- and FLV-treated cells, we still observed an average increase in IC as a function of IR similar to the untreated case (Fig. 3D), suggesting that the remaining intra-gene interactions observed after transcription inhibition may be attributed to residual Pol II occupancies. This ‘master curve’ provides further evidence that Pol II level only is predictive - in average - of the intra-gene folding whatever the conditions (treated or untreated).

**Fig 3.**
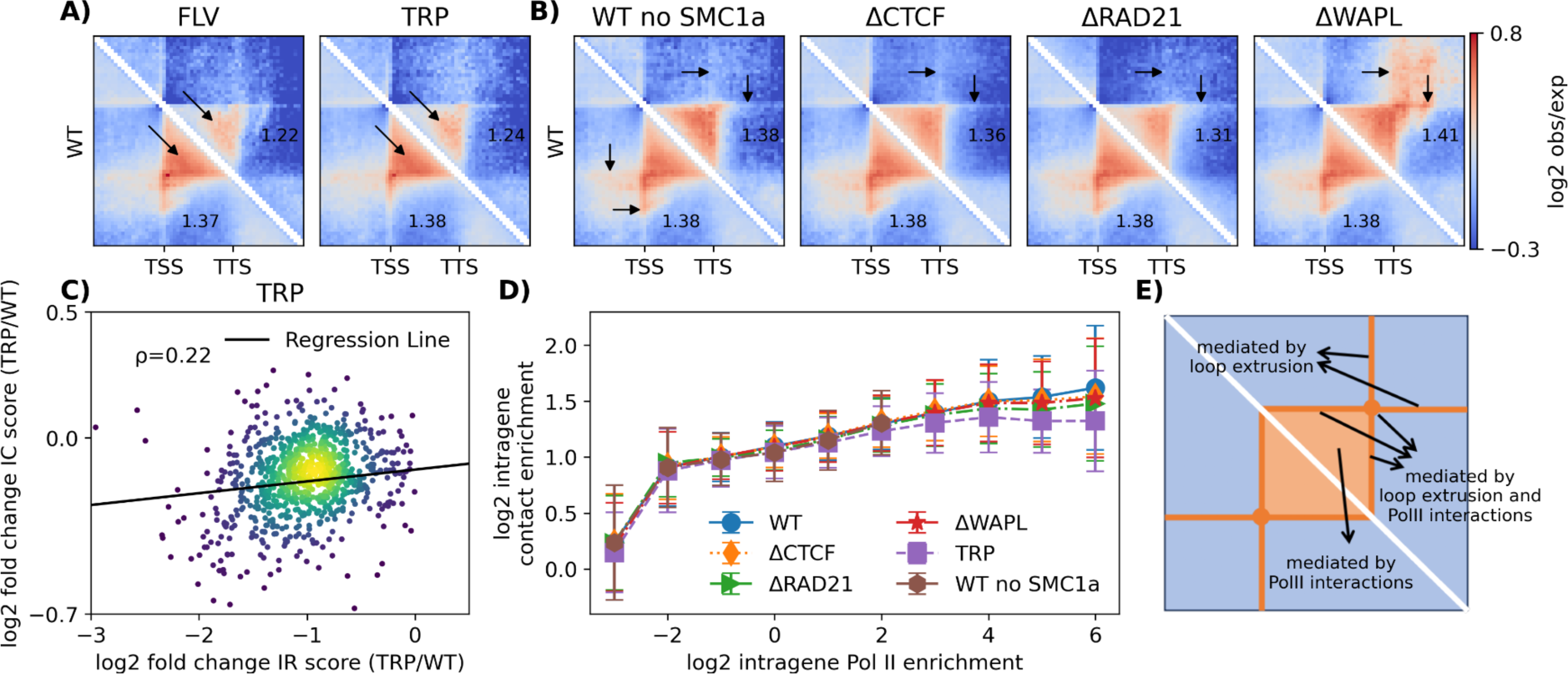
Gene conformation is affected by acute change in Pol II but not in CTCF and cohesin. (A) Comparison between PMGA of untreated WT cells and cells treated with transcription inhibitors for genes with size of 64-128 kb, IR_wt_>1 in WT condition and with a reduced IR score in treated cells (IR_treat_.<IR_wt_). (B) PMGA for genes with size of 64-128 kb and 1<IR_wt_<2 in conditions of reduced CTCF, RAD21 or WAPL levels or for a subset of genes with low SMC1a level in WT cells (WT no SMC1a, most left). (C) Scatter Plot of fold change of intragene contact enrichment against the fold change in Pol II occupancy after TRP treatment for the genes >64 kb, IR_wt_>1 in WT condition and with a reduced IR score in treated cells (IR_treat._<IR_wt_). The Spearman correlation is given. (D) The intragene contact enrichment upon acute depletion of RAD21, CTCF and WAPL, by IAA treatment of an engineered ES cell line, or by treatment with triptolide (TRP), as a function of IR score in the treated cells. The Spearman’s correlation between average IC and IR scores of WT is 0.97. The standard deviations of IC for each IR category are shown. (E) Scheme summarizing the different determinants of structures observed inside or around active genes.

In summary, our findings demonstrate a correlation between intra-gene condensation, interaction patterns, local transcriptional activity, gene length, and Pol II occupancy.

### Cohesin-mediated loop-extrusion activity plays a minor role on intra-gene condensation

Recent studies, both experimental and theoretical, have proposed that the loop extrusion mechanism, which plays a crucial role in the formation of TADs, might have an impact on the transcription machinery ^23,38,46,47,54^. Interestingly, we observed a significant correlation between the occupancy of CTCF and cohesin, the main players in loop extrusion, and the intra-gene contact enrichment and Pol II occupancy (Fig. 2C**,S1**). This observation led us to investigate whether cohesin-mediated loop extrusion could drive the correlation between transcriptional activity and intra-gene folding discussed earlier.

To address this question, we analyzed our original dataset from wild-type mESCs and excluded genes with high SMC1a (a cohesin subunit) occupancy (**Materials and methods**). The remaining genes, clustered based on IC and gene length, showed significantly lower levels of CTCF and cohesin, while Pol II profiles remained largely unchanged (**Fig.S13C**). Despite this subset of genes, the IC and IR scores still exhibited a strong correlation at a level similar to wild-type (Fig. 3D**,S13A,B**). Additionally, PMGA revealed that the typical interaction patterns observed within genes were still visible for cohesin-poor genes, although certain features such as stripes, which are known to be footprints of loop extrusion activity near extruding barriers ^38,46^, were absent outside the genes (black arrows in Fig. 3B **left**).

Furthermore, we utilized three publicly available mESC datasets where CTCF (ΔCTCF), the cohesin subunit RAD21 (ΔRAD21), or the cohesin unloader WAPL (ΔWAPL) were acutely depleted ^13^. These treatments led to significant alterations in CTCF and cohesin occupancies throughout the genome ^13^, as well as changes in TAD folding and CTCF-CTCF loops ^7,14^, such as a strong reduction in loop intensity in ΔCTCF and ΔRAD21 and reinforcement in ΔWAPL (**Fig.S8 bottom**). However, most gene expressions remained unaltered ^13^, and the majority of loops between Pol II peaks were unaffected (**Fig.S8 top**). Surprisingly, despite the acute changes in intra-gene CTCF and cohesin profiles (**Fig.S14C, S15C,S16C**), we observed only minimal effects on intra-gene interactions (Fig.3B**,D, Fig. S12,S14-16A,B**). The most noticeable - yet weak - changes in IC scores occurred in highly active genes (high IR), with an average 5% reduction in ΔRAD21 (Fig.3B). However, the changes in IC between WT and ΔRAD21 conditions did not exhibit a clear correlation with changes in RAD21 occupancy (**Fig.S17**). Similar to cohesin-poor genes in WT, the structural features associated with loop extrusion outside genes were lost or significantly reduced in ΔCTCF and ΔRAD21 (and enhanced in ΔWAPL) (Fig.3E).

Collectively, these results indicate that cohesin-mediated loop extrusion does not significantly affect the specific organization of transcribed genes, suggesting the presence of an independent mechanism.

### A biophysical model to investigate the role of transcription on gene folding

Our data analysis strongly suggests that Pol II occupancy drives the 3D organization of genes, independently of cohesin activity. Moreover, recent *in vitro* and *in vivo* experiments suggest that Pol IIs could form liquid-like droplets either directly through a phase-separation process mediated by weak interactions between their carboxy-terminal domains ^36,59–62^ or indirectly via the formation of Mediator condensates triggered by nascent RNAs ^63^. In the following, we developed a biophysical model to better characterize the phenomenology of Pol II-mediated gene folding by investigating how effective self-attractions between Pol II-occupied loci may shape the spatio-temporal dynamics of genes.

First, we built a stochastic model to describe Pol II occupancy and dynamics at a gene using a standard Totally Asymmetric Simple Exclusion Process (TASEP) ^64–66^. In this model (Fig. 4A, **Materials and methods**), Pol IIs can be loaded onto chromatin at the TSS with rate *α*, transcription elongation initiates with rate *γ*_0_, Pol IIs then progress along the gene at rate *γ* until they unbind from chromatin at TTS with a rate *β*. During this process, Pol IIs cannot overlap or bypass each other. We systematically varied the parameters of the TASEP model in order to predict different Pol II profiles along the gene at steady-state (**Fig. S18)**. For example, by varying model parameters (*γ*_0_/*γ* = 1, *β*/*γ* = 1 − *α*/*γ*), we reproduced uniform average profiles of Pol II occupancy along the gene, ranging from low (∼0.02) to high (∼0.80) densities (Fig. 4B**,C**).

**Fig. 4.**
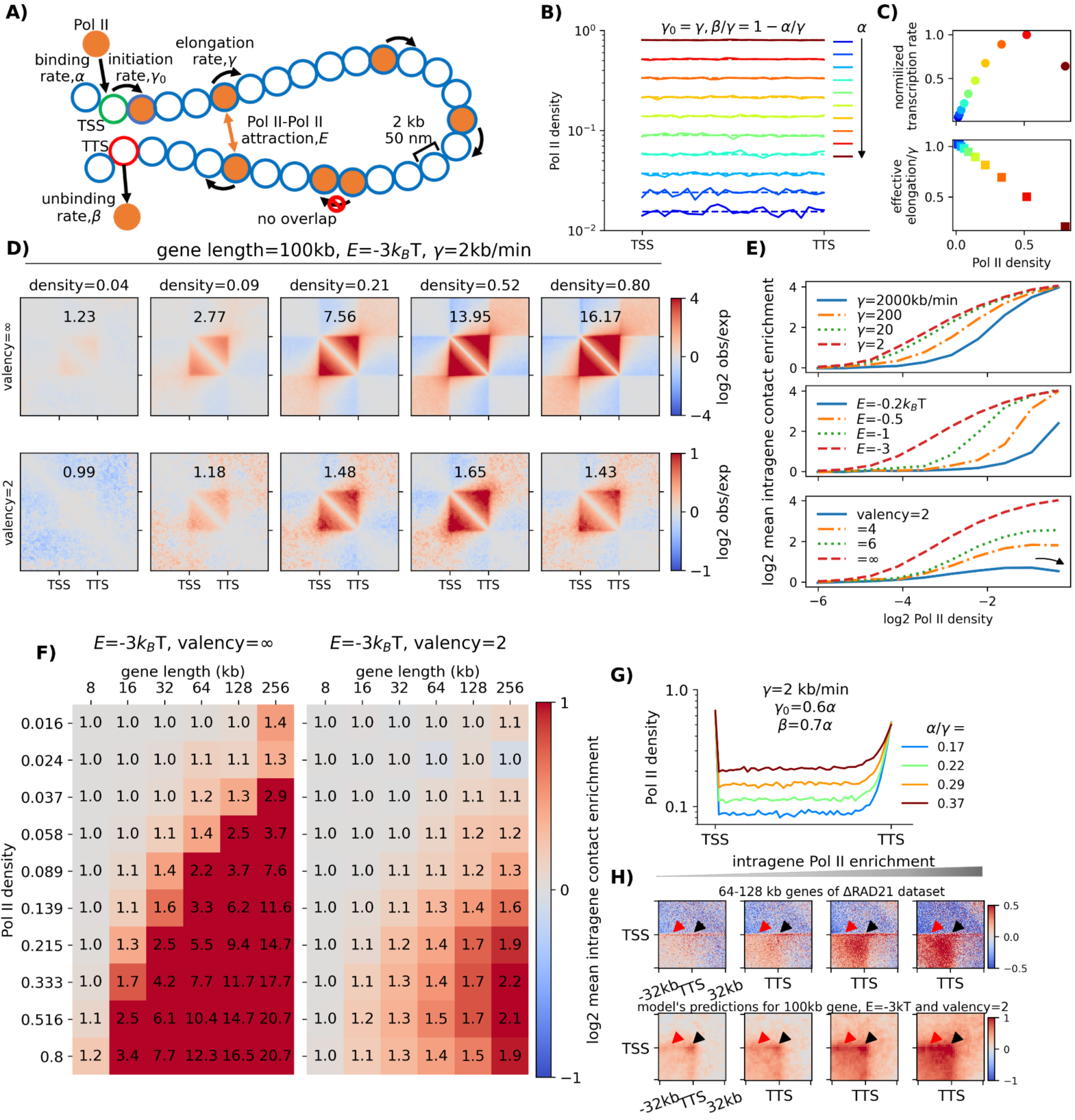
Transcription-mediated interactions regulate gene folding. (A) Schematic representation of the TASEP-decorated polymer model for gene transcription and 3D folding. (B) Pol II profiles along a 100kbp-long gene (L=50 monomers) for parameters tuned to generate a uniform occupancy along the gene, from low (blue) to high (red) densities. The solid and dashed curves are predictions from Monte-Carlo simulations and analytical calculations, respectively (see **Materials and Methods** and **Supplementary Notes**). (C) Normalized transcription rate (top), defined as the average number of Pol II unloadings from the TTS per time unit divided by its maximum, and normalized effective elongation rate (bottom), defined as the inverse of the time needed for one Pol II to fully transcribed a gene, as a function of Pol II density. (D) Predicted contact maps around a 100kbp-long gene for different Pol II densities and valencies. Corresponding IC scores are given. (E) IC versus IR curves as a function of the elongation rate *γ* (top), strength of interaction *E* (middle) and valency (bottom). (F) IC scores against IR scores (Pol II density) and gene length for two different valencies. The color bar is presented in a log2 scale, while the values are given in a linear scale. (G) Examples of non-uniform Pol II profiles having significant accumulations at TSS and TTS. (H) (Top) PMGA analysis of the contact around the TSS-TTS loop for 64-128 kb-long genes with increasing IR scores (from left to right) taken from ΔRAD21 dataset. (Bottom) Model predictions around the TSS-TTS loop for the non-uniform cases described in (G). TTS-TTS loops and promoter stripes are shown with black and red arrows, respectively.

Next, to assess the spatial organization of a gene, we integrated the TASEP in a 3D polymer model of chromatin fiber ^55,56^ (Fig. 4A, **Materials and methods**). Briefly, we represented a 20 Mbp-long section of chromatin as a self-avoiding chain (1 monomer = 2kbp = 50nm). We focused on a region of size *L* in the middle of the chain, which represents the gene of interest. Each monomer within the gene is characterized by a random binary variable indicating the local Pol II occupancy, whose dynamics is described by the TASEP. To investigate the impact of Pol IIs density and dynamics on gene folding, we assumed that monomers occupied by Pol II at a given time may self-interact at short-range with energy strength *E*. All the other monomers are considered non-interacting, neutral particles. The coupled stochastic spatio-temporal dynamics of the Pol II occupancies and 3D positions of the monomers are then simulated using kinetic Monte-Carlo (**Materials and methods**).

### Self-attraction between Pol II-bound genomic regions drives the intra-gene spatial organization

#### The coil-to-globule transition of a gene is controlled by Pol II occupancy and gene length

We quantified generic structural properties of the model and investigated the relationship between intra-gene condensation (IC scores) and Pol II density (IR scores) with respect to model parameters. In particular, we varied IR scores (via *α*/*γ*) while keeping other TASEP parameters constant, achieving uniform Pol II occupancies along the gene (as in Fig.4B**,C**), and we monitored the corresponding IC scores at steady-state.

For fixed gene length *L* and elongation rate *γ*, IC is an increasing function of both the Pol II density (Fig. 4D **upper panels**) and the strength *E* of self-attraction (Fig.4E **mid panel**): the gene’s polymeric subchain undergoes a theta-like collapse ^67^ towards a globular state when the Pol II occupancy reaches a critical value (**Movie S1**), such transition occurring at lower threshold densities for stronger interactions (*|E|*). Similar to standard self-interacting homopolymers ^68,69^, intra-gene contacts strengthen with increasing gene length (Fig.4F**, left panel**), while maintaining a fixed average Pol II level. This reflects the cooperative nature of the theta-collapse ^70,71^. At a constant average Pol II density, IC is a decreasing function of the Pol II elongation rate (Fig. 4E **upper panel**). Indeed, the capacity of Pol II-bound monomers to stably interact depends on the out-of-equilibrium dynamics of the elongating Pol IIs: shorter residence time of Pol II on a monomer (compared to typical polymer diffusion time) results in more transient Pol II-mediated interactions between monomers. Notably, biologically-relevant elongation rates (∼2kb/min, ^72^) correspond to the ‘slow’ elongation regime, maximizing gene condensation.

Overall, our model qualitatively recapitulates the global Pol II and gene length trends observed experimentally (Fig.2B). However, the predicted strengths of intra-gene contact enrichment are much stronger than expected (Fig. 4F **left panel**). For instance, a 128kbp-long gene shows a ∼6-fold increase in IC score with a ∼8-fold rise in Pol II density across the theta-collapse (for *γ* =2kb/min and *E*=-3 kT), whereas experimentally the same change in average Pol II occupancy yields only ∼35% increase in IC. We verified that reducing |*E*| does not resolve the problem as the theta-transition remains sharp and cooperative (**Fig. S19 left panel**).

#### Intra-gene condensation in mESC is consistent with a limited valency of Pol II-Pol II interactions

In our initial model, unrestricted interactions were allowed among the Pol II-occupied monomers in close proximity in the 3D space. However, such molecular interactions are mediated by only a restricted set of accessible residues and thus one monomer may have only a limited valency (number of simultaneous interactions).

Reducing the valency led to a global, sharp drop in intra-gene contact enrichment (Fig. 4D**, 4E lower panels, Movie S2**). For instance, at high Pol II occupancy, a ∼11-fold reduction in IC score was observed for valency 2 compared to unlimited valency. At lower valencies (2 or 3), the levels of contact enrichment aligned with experimental values (Fig. 4F **right panel, Fig. S19 right panel**) while still preserving the overall dependence on Pol II density and gene length see with unlimited valency.

However, an intriguing exception emerged: the IC score now displays a non-monotonic dependency with Pol II levels (Fig. 4E **lower panel**), as actually observed experimentally at high IR scores (Fig.1C). Within our framework, this behavior arises from a screening effect on long-range interactions. At high Pol II density, the neighboring Pol II-occupied monomers along the chain are likely to engage in interactions, limiting the ability of a monomer to interact with distantly located monomers and consequently reducing large-scale intra-gene condensation.

#### Nonuniform Pol II profiles lead to intra-gene architectural details

We previously focused on average gene folding properties by considering flat, homogeneous Pol II densities. However, experimental Pol II profiles show distinct peaks at TSS and TTS. By adjusting the TASEP parameters, we generated qualitatively similar peaked profiles of increasing density (Fig. 4G**, Materials and methods**). Using interacting parameters (*E=-3kT*, valency=2) compatible with the experimental IC vs IR relationship, we obtain for these nonuniform profiles very similar correlations between IC and IR scores and gene length (**Fig.S20**). Additionally, we predicted the formation of a stable loop between TSS and TTS in contact map as well as promoter-gene stripes within gene body, for high Pol II occupancy (Fig. 4H **lower panels**). Interestingly, off-diagonal pileup analysis of mESC Micro-C datasets around TSS-TTS anchors exhibits similar patterns independent of the cohesin loop-extrusion mechanism (Fig. 4H **upper panels, Fig.S21**), implying that such architectural details are driven by Pol II occupancy and effective Pol II-Pol II interactions.

### Modeling predicts a coupling between transcription and chromatin dynamics

#### Stochastic dynamics of gene folding in response to transcription bursting

Most mammalian genes undergo discontinuous transcription in bursts ^73–75^. To address the impact of such bursting kinetics on the gene spatio-temporal dynamics, we modified the TASEP model minimally: the promoter can stochastically switch between an “on” state, enabling Pol II binding and transcription, and an “off” state refractory to Pol II binding, with rates *k_on_* and *k_off_* (Fig. 5A). These rates define the effective Pol II binding rate (*α*_%*ff*_ = *αk_on_*/(*k_on_* + *k_off_*)), the burst frequency (= *k_on_k_off_*/(*k_on_* + *k_off_*), mean number of bursts per time unit) and the train size (= *α*/*k_off_*, mean number of Pol II binding and elongating during one burst). For simplicity, we assumed *k_on_* = *k_off_* ≡ *k*, allowing variation in burst properties from rare, long trains (*k=*0.01/min) to frequent, short ones (*k=*0.04/min) (Fig. 5B**,C**), while maintaining an almost constant average Pol II density profile (Fig. 5D).

**Fig. 5.**
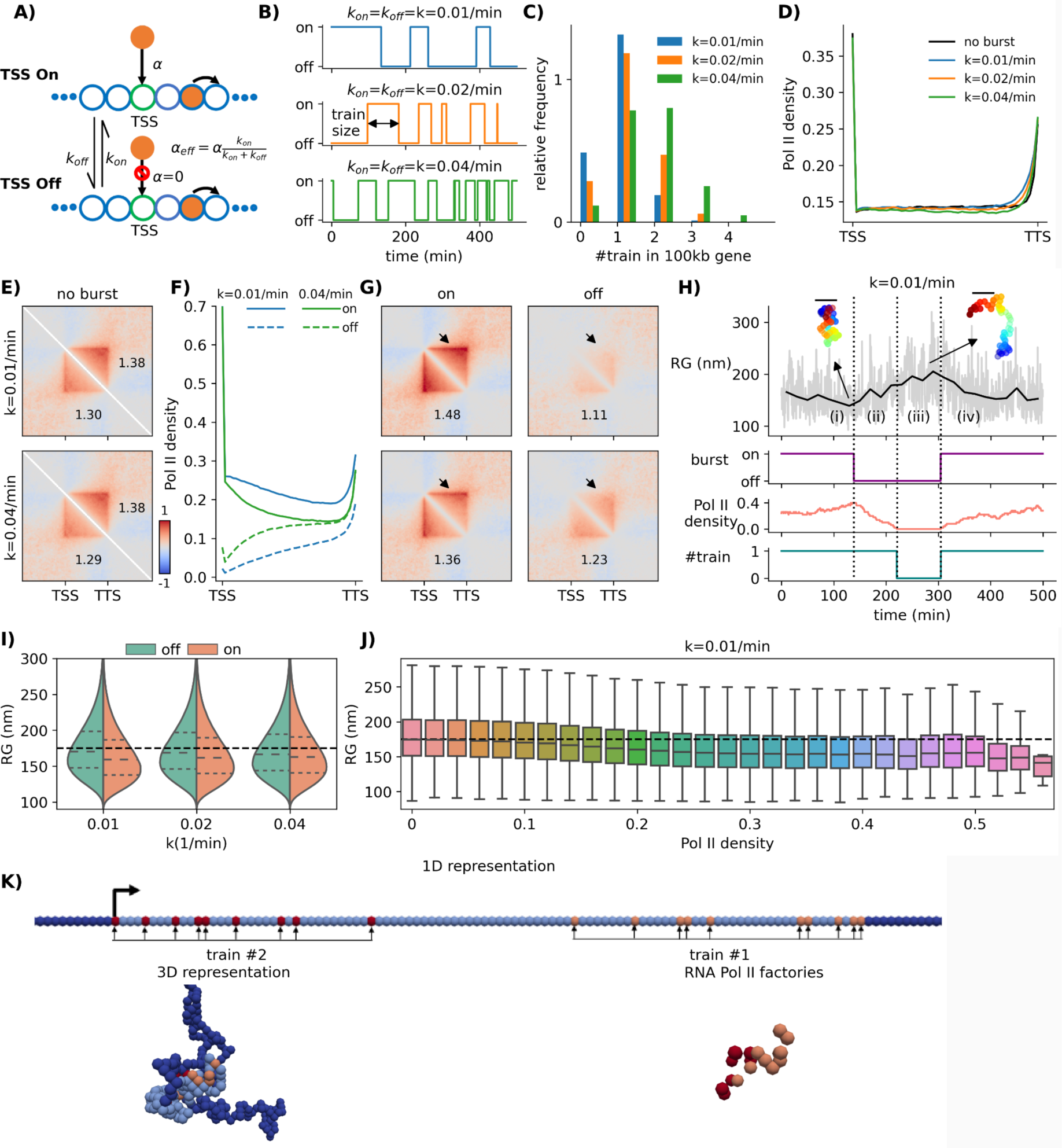
Transcriptional bursting leads to dynamical changes in gene conformation. (A) Schematic representation of transcriptional burst, where TSS alternatively switched on and off. (B) Three different examples of bursty gene activity ranging from long (k=0.01/min) to short (k=0.04/min) train size. (C) Probability distributions of the number of trains elongating on a gene at the same time for the three bursty regimes depicted in (B). (D) Average Pol II density profiles for the three bursty regimes depicted in (B) and in absence of burst. (E) Predicted contact maps with (lower left triangular part) and without burst (upper right triangular part) for long (top) and short (bottom) trains. (F) Pol II density profiles when TSS is “on” (solid lines) or “off” (dashed lines) for long (blue lines) and short (green lines) trains. (G) Predicted contact maps for conditions similar to (F). (H) (Top to bottom) Time evolution of the radius of gyration (RG) of a gene, TSS state, Pol II density along the gene and the number of trains elongating along the gene for k=0.01/min. Examples of 3D gene conformation are drawn when the gene is more or less condensed. Bars = 200nm. (I) Violin plots of RG in the “off” and “on” states for the three burst regimes in (B). The black dashed lines show the predictions for homopolymer model (i.e. zero interaction case). (J) Boxplot of RG as a function of the Pol II density for k=0.01/min. (K) A typical snapshot of gene 3D conformation (gene in light blue, flanking regions in dark blue) in the presence of two trains. The 1D representation shows the locations of Pol II-bound monomers for each train (orange and red dots). All simulations were done for a 100-kb gene with valency=2, E=-3 kT.

By averaging over all configurations, we observed a weak - but significant - decrease in intra-gene condensation in the presence of bursting (Fig. 5E). However, when considering the promoter’s on/off states separately, the impact of bursting became apparent with overall more intra-gene contacts and more pronounced TSS-TTS loops and promoter-gene stripes in the on-state (Fig. 5G). This effect was more pronounced for low burst frequency as the difference in Pol II occupancy between the on/off-states became more prominent (Fig. 5F). Similarly, more elongating trains lead to increased condensation (**Fig.S22**).

These findings suggest a time-correlation between transcriptional bursting and gene folding where dynamical changes in the gene’s radius of gyration (RG) are preceded by modifications in Pol II along the gene (Fig. 5H). Indeed, we observed an overall negative correlation between instantaneous Pol II density and RG, which was more pronounced for low burst frequencies (Fig. 5 **I,J**). Interestingly, when multiple trains are present simultaneously along the gene, the dynamic looping between could rise to the formation of ‘factories’ where they colocalize (Fig. 5K**, Movie S3**).

#### Transcription slows down gene mobility

Live-imaging experiments have indicated that chromatin motion is enhanced after Pol II inhibition or reduced after gene activation ^41,43,44^, suggesting a connection between transcription and a reduced gene mobility. To assess whether our biophysical model aligns with these observations, we computed for each monomer the mean-squared displacement (MSD), that measures the typical space explored by a locus over a time-lag Δt. We observed that *MSD*∼*D*Δ*t*^*δ*^, where *D* and *δ* are diffusion constant and exponent, respectively (Fig. 6A). *δ* ∼ 0.5 is independent of Pol II occupancy (Fig. 6B) and its value is consistent with live imaging experiments in mESC ^15,16^ and standard polymer dynamics ^76,77^. Conversely, *D* depends on Pol II density and gene length (Fig. 6C) with a perfect opposite trend as the intra-gene condensation (Fig. 4F **right**): the more condensed the gene the less mobile ^77^. For example, a 40-70% increase in intra-gene contacts corresponds to a 10-15% decrease for *D*, consistent with experiments (Fig. 6C**,D**).

**Fig. 6.**
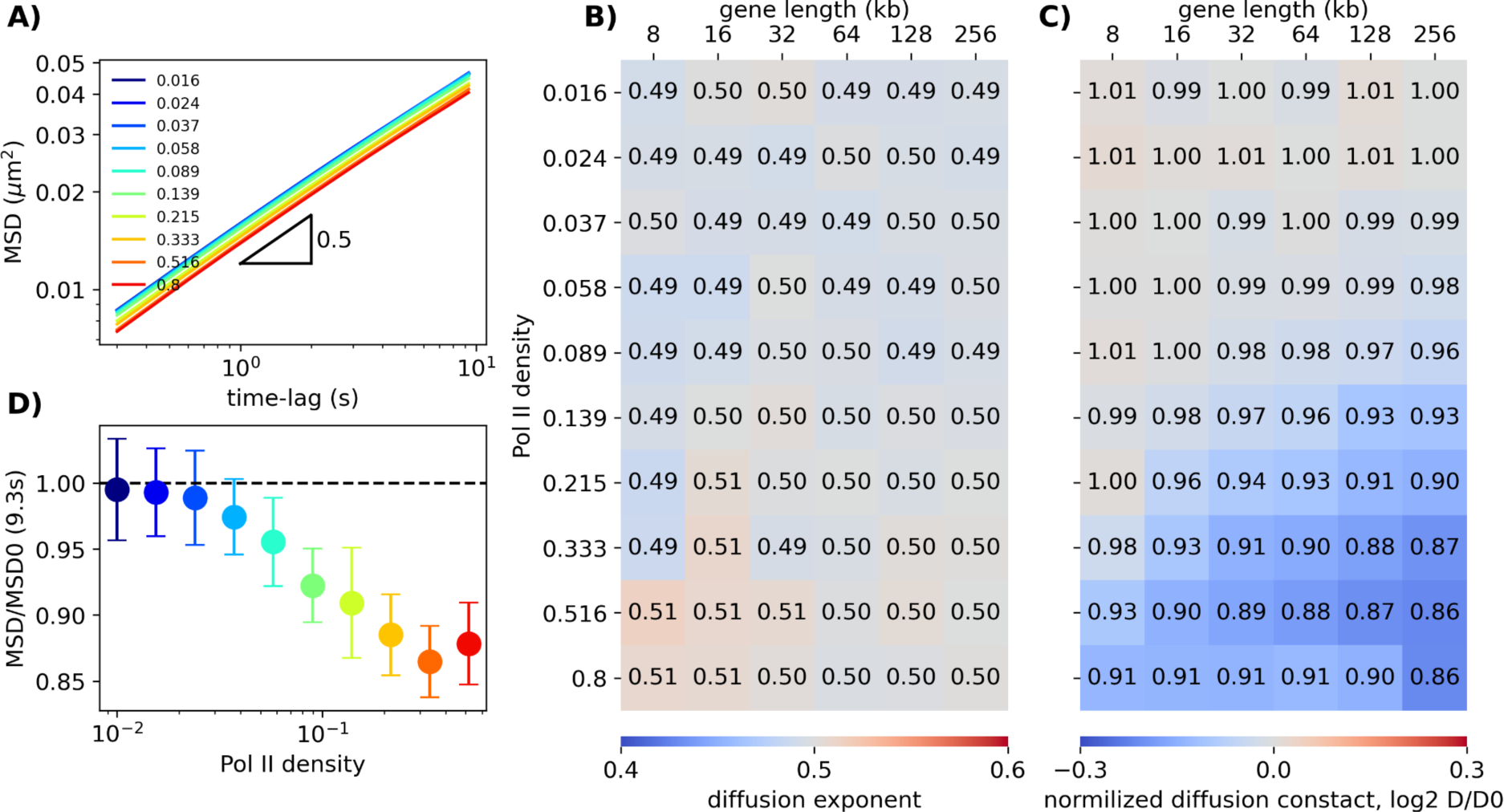
Transcription activity slows down gene mobility. (A) Mean-squared displacement *MSD*∼*D*Δ*t*^*δ*^ vs time-lag Δt for different Pol II densities for a 256kbp-long gene. (B) Diffusion exponent *δ* as a function of gene size and Pol II density. (C) As in (B) but for the diffusion constant *D* normalized by its value *D_0_* in absence of transcription. The color bar is presented in a log2 scale, while the values are given in a linear scale. (D) The ratio of MSD with (*MSD*) and without (*MSD0*) Pol II at t=9.3s as a function of Pol II density for a 256kbp-long gene. Colorscale as in (A). All simulations were done with valency=2, E=-3 kT.

### Transcription-associated long-range contacts between genes emerge from Pol II-mediated phase separation

Our analysis of intra-gene folding and dynamics suggests that similar mechanisms may explain the role of Pol II occupancy in distal inter-gene interactions. On the Micro-C map of mESC, we observed selective contact enrichments between distal highly active genes (Fig. 7**, Fig. S23**). For instance, the average contact frequency between the 811 kb-distant large active genes *Ahctf1* and *Parp1* is 3.2-fold higher than expected at similar genomic distance (Fig.7A). Both genes belong to the same A compartment, indicating that strongly transcribed genes may further colocalize within A. To test this hypothesis, we clustered all the 32-64 kb-long genes into three categories based on their IR score (Low, Mid and High) and performed PMGA (**Materials and methods**) of the inter-gene contacts for pairs of genes distant by more than 128 kb but less than 2 Mb (**Fig 7B, Fig.S24**). When both genes are transcribed (Mid and High clusters in **Fig 7B**), a strong promoter-promoter interaction is detected, as already observed in several studies ^13,27,29,78^. In addition, PMGA highlights that highly active genes (High-High) also exhibit significant contact enrichment between their gene bodies compared to the surrounding background in a transcription-dependent and loop extrusion-independent manner (**Fig.S24**). Contact enrichment between inactive genes (Low-Low) is similar to background and can be attributed to their location in the more compact B-compartment ^30^.

**Fig. 7.**
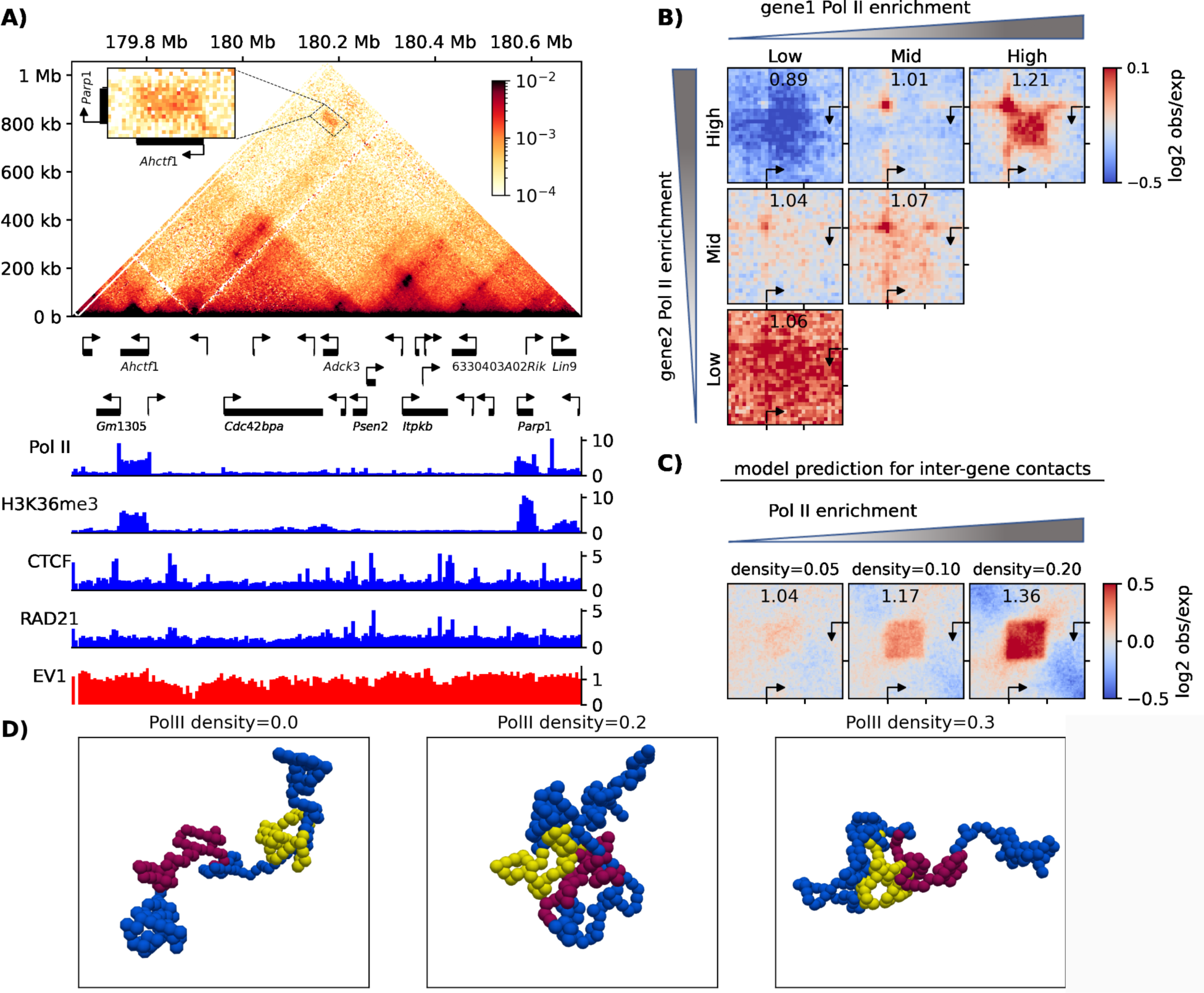
Inter-gene contacts between active genes. (A) Micro-C contact map of a ∼1 Mb region of mESC chromosome 1, with corresponding gene annotation and ChIP-seq profiles below. Inset shows a zoom between the long, highly-active genes of *Ahctf1* and *Parp1* (respectively, 58.7 kb and 32.3 kb-long and an expression of 22.4 FPKM and 151.4 FPKM). (B) Inter-gene pileup meta-gene analysis of the contact enrichment between two distant genes as a function of their intra-gene Pol II enrichment. (C) Model predictions for contacts between 60-kb-long genes for three different Pol II densities. (D) Examples of simulated 3D configurations illustrating the inter-gene interactions at various Pol II densities (gene regions in yellow and red, surrounding genomic regions in blue). All simulations were done with valency=2, E=-3 kT.

To rationalize these observations with our biophysical model, we conducted simulations for two 60 kbp-long genes distant by 460 kb, exhibiting similar steady-state Pol II profiles (Fig. 7C). We observed that Pol II-mediated interactions not only affect intra-gene contacts but also drive the formation of inter-gene contacts between TSS and TTS and between gene bodies, whose strengths increase with Pol II density. We obtained similar results for longer genes and shorter inter-gene distances (**Fig.S25**). Interestingly, interacting genes tend to colocalize and segregate from the rest of the simulated polymeric chain ^79,80^ (Fig.7D, **Movie S4**).

## Discussion

In this study, we analyzed publicly-available Micro-C data of mESC ^13,29^ to investigate the relationship between transcriptional activity and chromosome organization. Our findings align notably with previous studies ^29–31^. Specifically, we showed that, on average, at the single-gene level (2kbp-1Mbp), intra-gene contact enrichment, structural patterns (gene-loops, promoter-stripes, Fig.3E) and the degree of insulation from the surrounding genomic regions correlate positively with Pol II occupancy along the gene (Fig.1**,2**). Moreover, our study revealed that these observed features also exhibit a positive correlation with gene length (Fig.2), suggesting a cooperative mechanism for gene folding. Nevertheless, we noted a considerable degree of heterogeneity, implying that specific genes may deviate from the average behavior. These results stand in contrast with the very local structure of the chromatin fiber (< 600bp) that is increasingly open as transcription rate increases ^25^.

For highly expressed genes, we observed reduced contacts within gene body (Fig.1C), which aligns, although at a lesser extent, with the extended gene conformations observed for very long, highly expressed tissue-specific genes in mice ^37,39^.

Consistent with prior research ^47^, our results underscore the role of the loop extrusion process, recognized for driving the formation of loops and TADs ^11^ and reported to interfere with transcriptional elongation ^38,46,47^, into structural features outside the gene domain (Fig.3). However, in good agreement with recent high-precision Capture Micro-C data ^27^, we demonstrated that intragenic structure-function relation between gene condensation and gene transcription does not directly associate with loop extrusion ^79^ (Fig.3). At the inter-gene level, we observed long-range contacts between active genes, not only between gene promoters as already characterized ^81,82^, but also between gene bodies (Fig.7), here also closely tied to Pol II profiles and independent of loop extrusion (**Fig.S24**).

Altogether, our findings suggest that active genes are central units of the 3D genome ^25^ and form a subcompartment ^27,79,80^, driven by gene activity, Pol II binding and elongation. This observation likely holds true for other cell types, as we recently showed that intra-gene folding during mouse thymocyte maturation is, in average, also associated with change in transcription levels ^31^. The mechanisms described here are also likely to be broadly conserved in animals. Indeed, we analyzed the correlation between IC and IR (spearman’s ρ=0.48) in whole-embryo *Drosophila* data at embryonic nuclear cycle 14 (**Fig.S26**) ^83^. *Drosophila* is interesting as its chromosome organization is believed to be mainly driven by the spatial segregation of the epigenome instead of cohesin loop-extrusion processes ^4^. We found a similar nonmonotonic dependence of IC to IR as well as Pol II-related intra-gene interaction patterns. One exception is the effect of gene length that is less clear. Interestingly, in the bacterium *Escherichia coli*, higher transcription is also associated with more intra-gene contacts ^84^; in yeast and dinoflagellate, TAD-like structures are associated with (blocks of) active genes ^25,85^.

To better characterize the underlying mechanisms behind the correlations between Pol II activity and the transcriptionally-active subcompartment, we introduced a simple biophysical framework that accounts for the 1D dynamics of Pol II along genes coupled to the 3D polymer organization of chromosomes (Fig.4). Previous biophysical models have already addressed some aspects of Pol II-mediated phase separation via attractive ^35,48^ or active forces ^86^, focusing on large-scale inter-gene condensation, but never investigating intra-gene organization nor explicitly accounting for the transcription dynamics. By assuming self-attractive, short-range interactions between genomic loci bound to Pol II ^35,48^, our approach is able to recapitulate qualitatively the overall augmentation of intra-gene contacts associated to an enrichment of Pol II density inside gene body and to longer genes, consistent with a standard cooperative coil-globule transition observed for finite-size chains ^71,87–89^. Our model suggests that limiting the number of possible interactions per Pol II-bound region to low values (e.g., 2 or 3) allows to align quantitatively our predictions with experiments, leading to percolated but less condensed 3D domains ^90,91^. Interestingly, this constraint also explains the weak decompaction observed for highly transcribed genes as interactions between distant positions along the genes (mediating the large-scale gene’s condensation) are screened by (more frequent) interactions between nearest-neighbor Pol II-bound sites. This screening mechanism may also contribute to the formation of the extended transcription loops observed in long highly-transcribed genes ^37^, along with the potential stiffening of the chromatin fiber caused by the high density of nascent ribonucleoprotein complexes along the genes, as originally evoked.

Furthermore, our model predicts a strong coupling between gene structure and dynamics: transcription bursts may regulate the stochasticity of intra- and inter-gene contacts at the single-cell scale (Fig.5) ^92^; such dynamical contacts may conversely reduce locally gene mobility (Fig.6) and lead to long-range coherent motion between active regions ^56,93^, in good agreement with live-microscopy observations ^41,43,45^.

What are the molecular determinants of the putative attractive interaction between Pol II-bound loci hypothesized in our model? It is likely that several sources may directly or effectively participate in its regulation. The C-terminal domain (CTD) of Pol II can form liquid condensates *in vitro* under physiological conditions, which become unstable upon CTD phosphorylation ^59^. This mechanism may thus promote direct attractions *in vivo* between non-elongating Pol II, bound at promoters for example ^35^. CTDs may also interact with co-factors that can themselves phase-separate both at the transcriptional initiation ^34,94,95^ and elongation ^60,96^ stages, like FUS, BRD4, Mediator, P-TEFb or splicing factors. For example, the observed correlation between intra-gene condensation and the number of exons ^19^ at similar Pol II occupancy (**Fig.S4**) suggests a role for splicing-related condensates ^96^. In addition, transcription-generated supercoiling ^84,97^ or specific histone marks deposited along the gene bodies (that may regulate putative nucleosome-nucleosome interactions ^98^) may contribute to transcription-dependent effective interactions. The limited valency of interactions in our model aligns with a restricted number of simultaneously accessible residues involved in the aforementioned sources of Pol II-Pol II attraction. It is also possible that the screening effect observed at high transcription rates could be explained by the strength of interaction depending on local Pol II concentration and/or the length of nascent transcripts (**Supplementary Notes**), as RNA size and concentration can impact the stability of transcription-related condensates ^63^.

In conclusion, our results demonstrate the significant impact of Pol II binding and elongation on the spatiotemporal organization of the active genome through an out-of-equilibrium phase-separation process coupling the time-dependent dynamics of transcription to the formation of gene micro-domains and of transcriptionally-active subcompartment ^27,61,79,99^. This extends the concept of transcription factories ^100^, typically associated with inter-gene contacts, to the internal organization of long genes having multiple trains of transcribing Pol IIs. Consistent with our findings, recent works also proposed that interactions between Pol IIs may also facilitate promoter-enhancer communications ^23,101^. However, our approach provides only an “average” picture of the role of transcription on 3D chromosome organization and does not account for the various epigenetic, genomic and spatial factors that may interplay with Pol II-mediated phase separation ^47^ around specific genes, potentially explaining the variability of behaviors observed after transcription (de)activation ^31^.

Future investigations should aim to further elucidate the biological function(s) of such transcription-dependent micro-compartmentalization. Indeed, colocalization of active genomic regions may enhance the recycling of Pol II or transcription co-factors ^102,103^ by increasing their local concentrations. Investigating precisely such a “structure-function” coupling between the binding and assembly of transcription-associated components and condensates and the spatial folding of the genome remains an intriguing challenge and would require further developments both at the experimental and modeling levels.

## Materials and Methods

### Experimental data analysis

#### Datasets

The processed Micro-C data for mESCs (wild-type and mutants) and Drosophila in multi-resolution format *mcool* were downloaded from National Center for Biotechnology Information’s Gene Expression Omnibus (GEO) through accession no: GSE130275, GSE178982 and ArrayExpress accession E-MTAB-9306.

The ChIP-seq tracks, including Pol II, Pol II Ser5P and 2P, CTCF, RAD21, H3K27me3, H3K9me3 and H3K36me3, for wild type and different mutants in *BigWig* format were downloaded from GEO through accession no: GSE130275, GSE178982, GSE90893, GSE90994, GSE16013, GSE85191, GSE195830.

### Pileup meta-gene analysis (PMGA)

#### Contact maps

We used *cooltools* (https://github.com/open2c/cooltools)^104^ module to compute the obs/exp maps from the balanced contact maps, at various resolutions ranging from 100bp to 50kb. To perform intra-gene PMGA, for each gene *i* with size *l_i_* (>20 x resolution), we considered a domain of size 3*l_i_* around it, including the gene body and the two upstream and downstream flanking regions, each of size *l_i_*. To ensure consistency and facilitate pileup analysis, we rescaled each corresponding (3*l_i_*)*x*(3*l_i_*) obs/exp matrix to a (60,60)-pseudo-sized matrix by averaging the original matrix elements. An example of this rescaling process can be seen in **Figure S27**. We then aligned all the rescaled matrices in the transcription forward direction to maintain uniformity. Finally, we aggregated all the data of genes belonging to a given cluster (clustered by gene length, IR score, etc.).

For inter-gene PMGA, for each pair of genes, we considered the off-diagonal region of the obs/exp map of size (3*l*_1_)*x*(3*l*_2_) and centered at (*m*_1_, *m*_2_), with *l*_1_ and *m*_1_ the size and genomic position of the middle of gene 1 (same for gene 2). Then, similarly, we rescaled this region to a (30,30)-pseudo-matrix, aligned the genes in parallel forward direction and aggregated the pseudo-matrices belonging to the same cluster.

#### ChIP-seq tracks

Using *pyBigWig* (https://github.com/deeptools/pyBigWig), for each gene, we discretized the 3*l* domain (see above) into 60 bins and computed the coverage for each bin. Then, we aligned the domains in the transcription forward direction and aggregated over all genes in the same cluster.

### ChIP-seq peak calling and calculation of peak contacts

For each ChIP-seq track, we transformed *BigWig* to *bedGraph*, used MACS software ^105^ version 2 to call the peaks in the “no model” mode and merged the results from different replicates. Then, we sorted them by fold-change score (compared to input) and selected the most significant peaks (top 1/3). Finally, for every pair of peaks with a genomic distance between 160 and 320 kb, we used the off-diagonal pileup module of *cooltools* to compute the average peak contacts.

### Insulation score and compartments analysis

For computing the insulation score, we analyzed contact maps at 800-bp, 1600-bp and 3200-bp resolutions with the dedicated module of *cooltools* with sliding windows 3, 5, 10 and 25 times larger than the given resolution, e.g. 2.4, 4, 8 and 20-kb windows for 800-bp resolution. For the compartment analysis, we used the *eigs_cis* module of *cooltools* to compute the first eigenvector of the Pearson’s correlation matrix of contact map taking as inputs the 6.4-kbp resolution Micro-C maps and the GC coverage computing from *mm10* reference genome.

### Biophysical model

We previously introduced a self-avoiding semi-flexible polymer model for chromosomes ^55,56^. In this study, we employed a coarse-graining approach to represent a 20-Mbp-long chromatin fiber using 10,000 monomers. Each monomer corresponds to approximately 2-kbp of the genome and has a size of 50 nm (Fig. 4A). Within this chain, we inserted a 100-kbp-long gene (composed of 50 monomers), where TSS and TTS are located at the first and last monomers of the gene, respectively.

#### TASEP model

Each monomer *i* within a gene (of total size *n*) is characterized by a binary state *s_i_ε*{0,1} depending if a Pol II complex is bound to it (*s_i_* = 1) or not (*s_i_* = 0). We simulated the stochastic dynamics of Pol II binding, unbinding and elongation using a simple kinetic Monte-Carlo framework: each Monte Carlo step (MCS) consisted of (i) one attempt to bind a Pol II with rate *α* at the TSS if unoccupied (*s*_1_ = 0 → 1), (ii) one attempt to unbind Pol II with rate *β* at the TTS if occupied (*s_n_* = 1 → 0), and (iii) *n-1* elongation attempts, each consisting in randomly picking one monomer *i* in [1:n-1] and, if occupied, to move with rate *γ* the Pol II to its adjacent upstream monomer iff it is not already occupied ([*s_i_* = 1, *s_i_*_+1_ = 0] → [0,1]).

#### Polymer model

The polymer chain undergoes local movements on a FCC lattice with periodic boundary conditions under Metropolis criterion, as described in our previous works ^55^. The total Hamiltonian of a given configuration can be expressed as following:

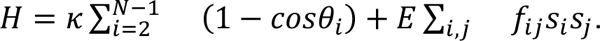

The first term accounts for the stiffness of the chain with *κ* the bending rigidity and *θ_i_* the local bending angle at monomer *i*. The second term represents the Pol II-Pol II interaction, where *E* denotes the attractive interaction strength, and *f_ij_* equals 1 if monomers *i* and *j* occupy nearest neighboring sites on the lattice.

For simulations with a limited valency number, we defined an interaction list for each monomer with *s_i_* = 1. This list stores the genomic positions of the other monomers it interacts with and is constrained not to exceed the given valency number. It is updated after any polymer or TASEP moves.

Note that, due to the relatively high stall force of Pol II (∼25-30 pN ^106^), we assumed that Pol II-Pol II interactions do not impede Pol II elongation.

#### Numerical simulations

In our study, we set *κ* ∼ 1.2 *k_B_T* and a lattice volumic density of 50% to account for a chromatin fiber with a Kuhn length of 100 nm ^69,107^ and a typical base-pair density found in mammalian and fly genomes (∼0.01 bp/nm³) ^108^. Simulations were initiated by unknotted configurations ^55^ and performed with a kinetic Monte Carlo algorithm. In addition to the TASEP moves (see above), each MCS contains N local polymer trial moves. For each parameter set, 20 independent trajectories were conducted, discarding the first 10⁶ MCS from each trajectory to allow the system to reach a steady state. Subsequently, snapshots of the system were saved every 10³ MCS during the simulation during 10⁷ MCS and analyzed subsequently (see below). Since the characteristic spatial and time scales of the phenomenon under study (∼100s nm, ∼min) are well beyond the discretization scales (50 nm, 3 msec) imposed by the lattice and the kinetic Monte-Carlo algorithm, the obtained results are not expected to depend qualitatively on the underlying modeling and simulation frameworks ^109^.

#### Data analysis

The radius of gyration (RG) provides a measure of the typical spatial extent of a gene, reflecting its overall span in 3D space. In a given configuration, the position of monomer *i* can be defined as 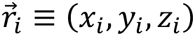. The RG is then calculated as follows:

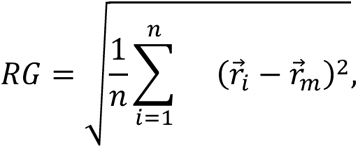

where 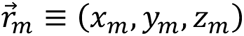 is the mean position of all monomers.

To extract the diffusion coefficient (*D_i_*) and exponent (*α_i_*) for monomer *i*, we first computed the time-averaged and ensemble-averaged mean-squared displacement, < *MSD_i_* >, as a function of the time-lag, △ *t*. Subsequently, we performed a power-law fit of the form *D_i_* △ *t^αi^* to the resulting curve using Numpy function *numpy.polyfit(log*△ *t,log*< *MSD_i_* >*,1)*.

Furthermore, to establish a correspondence between simulation (MCS) and real (seconds) times, we compared our predictions with the typical MSD observed in yeast (∼0.01(*μm*^2^/*sec*^0.5^) △ *t*^0.5^, with △ *t* in seconds) ^76^, leading to 1 MCS∼3 msec.

#### Data and code availability

Processed data (intra-gene contact, RNA-seq and ChIP-seq enrichments, compartments and exon numbers for each gene > 1kbp), Python notebooks for PMGA analysis and simulation codes are available on GitHub (https://github.com/physical-biology-of-chromatin/Transcription).

## Acknowledgment

We are grateful to Xavier Darzacq’s lab for sharing processed data; Marco Di Stefano, Guillermo Orsi and Aurèle Piazza for critical reading of the manuscript; Kerstin Bystricky, Tom Sexton, Giacomo Cavalli, Cédric Vaillant and the members of the Jost lab for fruitful discussions. We acknowledge Agence Nationale de la Recherche [ANR-18-CE12-0006-03, ANR-18-CE45-0022-01, ANR-21-CE45-0011-01] for funding. We thank PSMN (Pôle Scientifique de Modélisation Numérique) of the ENS de Lyon for computing resources.

## Author contributions

H.S. and D.J. designed the research; D.J. supervised the project; H.S. developed analytical tools and performed the research; H.S. and D.J. analyzed the data; G.F. provided conceptual advice; H.S. and D.J. wrote the paper with input from G.F.

## Reference

1. Eagen, K. P. Principles of Chromosome Architecture Revealed by Hi-C. Trends Biochem. Sci. 43, 469–478 (2018).

2. Jerkovic, I. & Cavalli, G. Understanding 3D genome organization by multidisciplinary methods. Nat. Rev. Mol. Cell Biol. 22, 511–528 (2021).

3. Lieberman-Aiden, E. et al. Comprehensive mapping of long-range interactions reveals folding principles of the human genome. Science 326, 289–293 (2009).

4. Rowley, M. J. et al. Evolutionarily Conserved Principles Predict 3D Chromatin Organization. Mol. Cell 67, 837–852.e7 (2017).

5. Dixon, J. R. et al. Topological domains in mammalian genomes identified by analysis of chromatin interactions. Nature 485, 376–380 (2012).

6. Nora, E. P. et al. Spatial partitioning of the regulatory landscape of the X-inactivation centre. Nature 485, 381–385 (2012).

7. Rao, S. S. P. et al. Cohesin Loss Eliminates All Loop Domains. Cell 171, 305–320.e24 (2017).

8. Mirny, L. A., Imakaev, M. & Abdennur, N. Two major mechanisms of chromosome organization. Curr. Opin. Cell Biol. 58, 142–152 (2019).

9. Wang, L. et al. Histone Modifications Regulate Chromatin Compartmentalization by Contributing to a Phase Separation Mechanism. Mol. Cell 76, 646–659.e6 (2019).

10. Zenk, F. et al. HP1 drives de novo 3D genome reorganization in early Drosophila embryos. Nature 593, 289–293 (2021).

11. Fudenberg, G. et al. Formation of Chromosomal Domains by Loop Extrusion. Cell Reports 15, 2038–2049 (2016).

12. Guo, Y. et al. CRISPR Inversion of CTCF Sites Alters Genome Topology and Enhancer/Promoter Function. Cell 162, 900–910 (2015).

13. Hsieh, T.-H. S. et al. Enhancer–promoter interactions and transcription are largely maintained upon acute loss of CTCF, cohesin, WAPL or YY1. Nat. Genet. 54, 1919–1932 (2022).

14. Schwarzer, W. et al. Two independent modes of chromatin organization revealed by cohesin removal. Nature 551, 51–56 (2017).

15. Gabriele, M. et al. Dynamics of CTCF- and cohesin-mediated chromatin looping revealed by live-cell imaging. Science 376, 496–501 (2022).

16. Mach, P. et al. Cohesin and CTCF control the dynamics of chromosome folding. Nat. Genet. 54, 1907–1918 (2022).

17. Wutz, G. et al. Topologically associating domains and chromatin loops depend on cohesin and are regulated by CTCF, WAPL, and PDS5 proteins. EMBO J. 36, 3573–3599 (2017).

18. Lupiáñez, D. G. et al. Disruptions of topological chromatin domains cause pathogenic rewiring of gene-enhancer interactions. Cell 161, 1012–1025 (2015).

19. Bonev, B. et al. Multiscale 3D Genome Rewiring during Mouse Neural Development. Cell 171, 557–572.e24 (2017).

20. Zuin, J. et al. Nonlinear control of transcription through enhancer-promoter interactions. Nature 604, 571–577 (2022).

21. Chen, H. et al. Dynamic interplay between enhancer-promoter topology and gene activity. Nat. Genet. 50, 1296–1303 (2018).

22. Barshad, G. et al. RNA polymerase II dynamics shape enhancer-promoter interactions. Nat. Genet. 55, 1370–1380 (2023).

23. Zhang, S., Übelmesser, N., Barbieri, M. & Papantonis, A. Enhancer-promoter contact formation requires RNAPII and antagonizes loop extrusion. Nat. Genet. 55, 832–840 (2023).

24. Hilbert, L. et al. Transcription organizes euchromatin via microphase separation. Nat. Commun. 12, 1360 (2021).

25. Hsieh, T.-H. S. et al. Mapping Nucleosome Resolution Chromosome Folding in Yeast by Micro-C. Cell 162, 108–119 (2015).

26. Aljahani, A. et al. Analysis of sub-kilobase chromatin topology reveals nano-scale regulatory interactions with variable dependence on cohesin and CTCF. Nat. Commun. 13, 2139 (2022).

27. Goel, V. Y., Huseyin, M. K. & Hansen, A. S. Region Capture Micro-C reveals coalescence of enhancers and promoters into nested microcompartments. Nat. Genet. 55, 1048–1059 (2023).

28. Balasubramanian, D. et al. Enhancer-promoter interactions can form independently of genomic distance and be functional across TAD boundaries. Nucleic Acids Res. gkad1183 (2023).

29. Hsieh, T.-H. S. et al. Resolving the 3D Landscape of Transcription-Linked Mammalian Chromatin Folding. Mol. Cell 78, 539–553.e8 (2020).

30. Rowley, M. J. et al. Condensin II Counteracts Cohesin and RNA Polymerase II in the Establishment of 3D Chromatin Organization. Cell Rep. 26, 2890–2903.e3 (2019).

31. Chahar, S., Zouari, Y. B., Salari, H., Molitor, A. M. & Kobi, D. Context-dependent transcriptional remodeling of TADs during differentiation. PLoS Biol. 21, e3002424 (2023).

32. Li, G. et al. Extensive promoter-centered chromatin interactions provide a topological basis for transcription regulation. Cell 148, 84–98 (2012).

33. Cisse, I. I. et al. Real-time dynamics of RNA polymerase II clustering in live human cells. Science 341, 664–667 (2013).

34. Cho, W.-K. et al. Mediator and RNA polymerase II clusters associate in transcription-dependent condensates. Science 361, 412–415 (2018).

35. Pancholi, A. et al. RNA polymerase II clusters form in line with surface condensation on regulatory chromatin. Mol. Syst. Biol. 17, e10272 (2021).

36. Hnisz, D., Shrinivas, K., Young, R. A., Chakraborty, A. K. & Sharp, P. A. A Phase Separation Model for Transcriptional Control. Cell 169, 13–23 (2017).

37. Leidescher, S. et al. Spatial organization of transcribed eukaryotic genes. Nat. Cell Biol. 24, 327–339 (2022).

38. Heinz, S. et al. Transcription Elongation Can Affect Genome 3D Structure. Cell 174, 1522– 1536.e22 (2018).

39. Winick-Ng, W. et al. Cell-type specialization is encoded by specific chromatin topologies. Nature 599, 684–691 (2021).

40. Jiang, Y. et al. Genome-wide analyses of chromatin interactions after the loss of Pol I, Pol II, and Pol III. Genome Biol. 21, 158 (2020).

41. Germier, T. et al. Real-Time Imaging of a Single Gene Reveals Transcription-Initiated Local Confinement. Biophys. J. 113, 1383–1394 (2017).

42. Gu, B. et al. Transcription-coupled changes in nuclear mobility of mammalian cis-regulatory elements. Science 359, 1050–1055 (2018).

43. Nagashima, R. et al. Single nucleosome imaging reveals loose genome chromatin networks via active RNA polymerase II. Journal of Cell Biology 218, 1511–1530 (2019).

44. Shaban, H. A., Barth, R., Recoules, L. & Bystricky, K. Hi-D: nanoscale mapping of nuclear dynamics in single living cells. Genome Biol. 21, 95 (2020).

45. Barth, R. & Shaban, H. A. Spatially coherent diffusion of human RNA Pol II depends on transcriptional state rather than chromatin motion. Nucleus 13, 194–202 (2022).

46. Brandão, H. B. et al. RNA polymerases as moving barriers to condensin loop extrusion. Proc. Natl. Acad. Sci. U. S. A. 116, 20489–20499 (2019).

47. Banigan, E. J. et al. Transcription shapes 3D chromatin organization by interacting with loop extrusion. Proc. Natl. Acad. Sci. U. S. A. 120, e2210480120 (2023).

48. Cook, P. R. & Marenduzzo, D. Transcription-driven genome organization: a model for chromosome structure and the regulation of gene expression tested through simulations. Nucleic Acids Res. 46, 9895–9906 (2018).

49. Larkin, J. D., Papantonis, A., Cook, P. R. & Marenduzzo, D. Space exploration by the promoter of a long human gene during one transcription cycle. Nucleic Acids Res. 41, 2216–2227 (2013).

50. Lengronne, A. et al. Cohesin relocation from sites of chromosomal loading to places of convergent transcription. Nature 430, 573–578 (2004).

51. Busslinger, G. A. et al. Cohesin is positioned in mammalian genomes by transcription, CTCF and Wapl. Nature 544, 503–507 (2017).

52. Valton, A.-L. et al. A cohesin traffic pattern genetically linked to gene regulation. Nat. Struct. Mol. Biol. 29, 1239–1251 (2022).

53. Zhang, S. et al. RNA polymerase II is required for spatial chromatin reorganization following exit from mitosis. Sci Adv 7, eabg8205 (2021).

54. Rivosecchi, J. et al. RNA polymerase backtracking results in the accumulation of fission yeast condensin at active genes. Life Sci Alliance 4, e202101046 (2021).

55. Ghosh, S. K. & Jost, D. How epigenome drives chromatin folding and dynamics, insights from efficient coarse-grained models of chromosomes. PLoS Comput. Biol. 14, e1006159 (2018).

56. Salari, H., Di Stefano, M. & Jost, D. Spatial organization of chromosomes leads to heterogeneous chromatin motion and drives the liquid- or gel-like dynamical behavior of chromatin. Genome Res. 32, 28–43 (2022).

57. Bartkowiak, B. & Greenleaf, A. L. Phosphorylation of RNAPII: To P-TEFb or not to P-TEFb? Transcription 2, 115–119 (2011).

58. Belaghzal, H. et al. Liquid chromatin Hi-C characterizes compartment-dependent chromatin interaction dynamics. Nat. Genet. 53, 367–378 (2021).

59. Boehning, M. et al. RNA polymerase II clustering through carboxy-terminal domain phase separation. Nat. Struct. Mol. Biol. 25, 833–840 (2018).

60. Lu, H. et al. Phase-separation mechanism for C-terminal hyperphosphorylation of RNA polymerase II. Nature 558, 318–323 (2018).

61. Rippe, K. & Papantonis, A. Functional organization of RNA polymerase II in nuclear subcompartments. Curr. Opin. Cell Biol. 74, 88–96 (2022).

62. Phatnani, H. P. & Greenleaf, A. L. Phosphorylation and functions of the RNA polymerase II CTD. Genes Dev. 20, 2922–2936 (2006).

63. Henninger, J. E., et al. RNA-Mediated Feedback Control of Transcriptional Condensates. Cell vol. 184 207–225.e24 Preprint at 10.1016/j.cell.2020.11.030 (2021).

64. Bressloff, P. C. & Newby, J. M. Stochastic models of intracellular transport. Rev. Mod. Phys. 85, 135–196 (2013).

65. Schadschneider, A., Chowdhury, D. & Nishinari, K. Stochastic Transport in Complex Systems: From Molecules to Vehicles. (Elsevier, 2010).

66. Mines, R. C., Lipniacki, T. & Shen, X. Slow nucleosome dynamics set the transcriptional speed limit and induce RNA polymerase II traffic jams and bursts. PLoS Comput. Biol. 18, e1009811 (2022).

67. de Gennes, P.-G. & Gennes, P.-G. Scaling Concepts in Polymer Physics. (Cornell University Press, 1979).

68. Lesage, A., Dahirel, V., Victor, J.-M. & Barbi, M. Polymer coil–globule phase transition is a universal folding principle of Drosophila epigenetic domains. Epigenetics Chromatin 12, 28 (2019).

69. Socol, M. et al. Rouse model with transient intramolecular contacts on a timescale of seconds recapitulates folding and fluctuation of yeast chromosomes. Nucleic Acids Res. 47, 6195–6207 (2019).

70. Grassberger, P. & Hegger, R. Simulations of three-dimensional θ polymers. J. Chem. Phys. 102, 6881–6899 (1995).

71. Caré, B. R., Carrivain, P., Forné, T., Victor, J.-M. & Lesne, A. Finite-Size Conformational Transitions: A Unifying Concept Underlying Chromosome Dynamics. Commun. Theor. Phys. 62, 607 (2014).

72. Jonkers, I., Kwak, H. & Lis, J. T. Genome-wide dynamics of Pol II elongation and its interplay with promoter proximal pausing, chromatin, and exons. Elife 3, e02407 (2014).

73. Fukaya, T., Lim, B. & Levine, M. Enhancer Control of Transcriptional Bursting. Cell 166, 358–368 (2016).

74. Tunnacliffe, E. & Chubb, J. R. What is a transcriptional burst? Trends Genet. 36, 288–297 (2020).

75. Dar, R. D. et al. Transcriptional burst frequency and burst size are equally modulated across the human genome. Proc. Natl. Acad. Sci. U. S. A. 109, 17454–17459 (2012).

76. Hajjoul, H. et al. High-throughput chromatin motion tracking in living yeast reveals the flexibility of the fiber throughout the genome. Genome Res. 23, 1829–1838 (2013).

77. Tortora, M. M., Salari, H. & Jost, D. Chromosome dynamics during interphase: a biophysical perspective. Curr. Opin. Genet. Dev. 61, 37–43 (2020).

78. Schoenfelder, S. et al. The pluripotent regulatory circuitry connecting promoters to their long-range interacting elements. Genome Res. 25, 582–597 (2015).

79. Miron, E. et al. Chromatin arranges in chains of mesoscale domains with nanoscale functional topography independent of cohesin. Sci Adv 6, (2020).

80. Gelléri, M. et al. True-to-scale DNA-density maps correlate with major accessibility differences between active and inactive chromatin. Cell Rep. 42, 112567 (2023).

81. Joshi, O. et al. Dynamic Reorganization of Extremely Long-Range Promoter-Promoter Interactions between Two States of Pluripotency. Cell Stem Cell 17, 748–757 (2015).

82. Zhao, L. et al. Chromatin loops associated with active genes and heterochromatin shape rice genome architecture for transcriptional regulation. Nat. Commun. 10, 3640 (2019).

83. Ing-Simmons, E. et al. Independence of chromatin conformation and gene regulation during Drosophila dorsoventral patterning. Nat. Genet. 53, 487–499 (2021).

84. Bignaud, A. et al. Transcriptional units form the elementary constraining building blocks of the bacterial chromosome. bioRxiv 2022.09.16.507559 (2022) doi:10.1101/2022.09.16.507559.

85. Nand, A. et al. Genetic and spatial organization of the unusual chromosomes of the dinoflagellate Symbiodinium microadriaticum. Nat. Genet. 53, 618–629 (2021).

86. Shin, S., Shi, G., Cho, H. W. & Thirumalai, D. Transcription-induced active forces suppress chromatin motion. bioRxiv (2022) doi:10.1101/2022.04.30.490180.

87. Abdulla, A. Z., Tortora, M. M. C., Vaillant, C. & Jost, D. Topological constraints and finite-size effects in quantitative polymer models of chromatin organization. Macromolecules 56, 8697–8709 (2023).

88. Conte, M. et al. Dynamic and equilibrium properties of finite-size polymer models of chromosome folding. Phys Rev E 104, 054402 (2021).

89. R. Caré, B., Emeriau, P.-E., Cortini, R. & Victor, J.-M. Chromatin epigenomic domain folding: size matters. AIMS Biophys. 2, 517–530 (2015).

90. Ryu, J.-K. et al. Bridging-induced phase separation induced by cohesin SMC protein complexes. Sci Adv 7, (2021).

91. Zeng, X. & Pappu, R. V. Developments in describing equilibrium phase transitions of multivalent associative macromolecules. Curr. Opin. Struct. Biol. 79, 102540 (2023).

92. Giorgetti, L. et al. Predictive polymer modeling reveals coupled fluctuations in chromosome conformation and transcription. Cell 157, 950–963 (2014).

93. Di Stefano, M. et al. Transcriptional activation during cell reprogramming correlates with the formation of 3D open chromatin hubs. Nat. Commun. 11, 2564 (2020).

94. Sabari, B. R. et al. Coactivator condensation at super-enhancers links phase separation and gene control. Science 361, (2018).

95. Kwon, I. et al. Phosphorylation-regulated binding of RNA polymerase II to fibrous polymers of low-complexity domains. Cell 155, 1049–1060 (2013).

96. Guo, Y. E. et al. Pol II phosphorylation regulates a switch between transcriptional and splicing condensates. Nature 572, 543–548 (2019).

97. Portman, J. R., Brouwer, G. M., Bollins, J., Savery, N. J. & Strick, T. R. Cotranscriptional R-loop formation by Mfd involves topological partitioning of DNA. Proc. Natl. Acad. Sci. U. S. A. 118, (2021).

98. Gibson, B. A. et al. Organization of Chromatin by Intrinsic and Regulated Phase Separation. Cell 179, 470–484.e21 (2019).

99. Nozaki, T. et al. Condensed but liquid-like domain organization of active chromatin regions in living human cells. Sci Adv 9, eadf1488 (2023).

100. Cook, P. R. A model for all genomes: the role of transcription factories. J. Mol. Biol. 395, 1– 10 (2010).

101. Barshad, G. et al. RNA polymerase II and PARP1 shape enhancer-promoter contacts. Nat. Genet. 55, 1370–1380 (2023).

102. Chiang, M. et al. Gene structure heterogeneity drives transcription noise within human chromosomes. bioRxiv 2022.06.09.495447 (2022) doi:10.1101/2022.06.09.495447.

103. Semeraro, M., et al. A multicolour polymer model for the prediction of 3D structure and transcription in human chromatin. bioRxiv (2023) doi:10.1101/2023.01.16.524198.

104. Open2C et al. Cooltools: enabling high-resolution Hi-C analysis in Python. bioRxiv 2022.10.31.514564 (2022) doi:10.1101/2022.10.31.514564.

105. Zhang, Y. et al. Model-based analysis of ChIP-Seq (MACS). Genome Biol. 9, R137 (2008).

106. Wang, H. Y., Elston, T., Mogilner, A. & Oster, G. Force generation in RNA polymerase. Biophys. J. 74, 1186–1202 (1998).

107. Arbona, J.-M., Herbert, S., Fabre, E. & Zimmer, C. Inferring the physical properties of yeast chromatin through Bayesian analysis of whole nucleus simulations. Genome Biol. 18, 81 (2017).

108. Milo, R., Jorgensen, P., Moran, U., Weber, G. & Springer, M. BioNumbers—the database of key numbers in molecular and cell biology. Nucleic Acids Res. 38, D750–D753 (2009).

109. Halverson, J. D., Kremer, K. & Grosberg, A. Y. Comparing the results of lattice and off-lattice simulations for the melt of nonconcatenated rings. J. Phys. A: Math. Theor. 46, 065002 (2013).

