## Supplementary Materials for "Transcription regulates the spatio-temporal dynamics of genes through micro-compartmentalization"

**This PDF file includes:**

Supplementary figures S1 to S27

Supplementary Notes

Legends for Movies S1 to S4

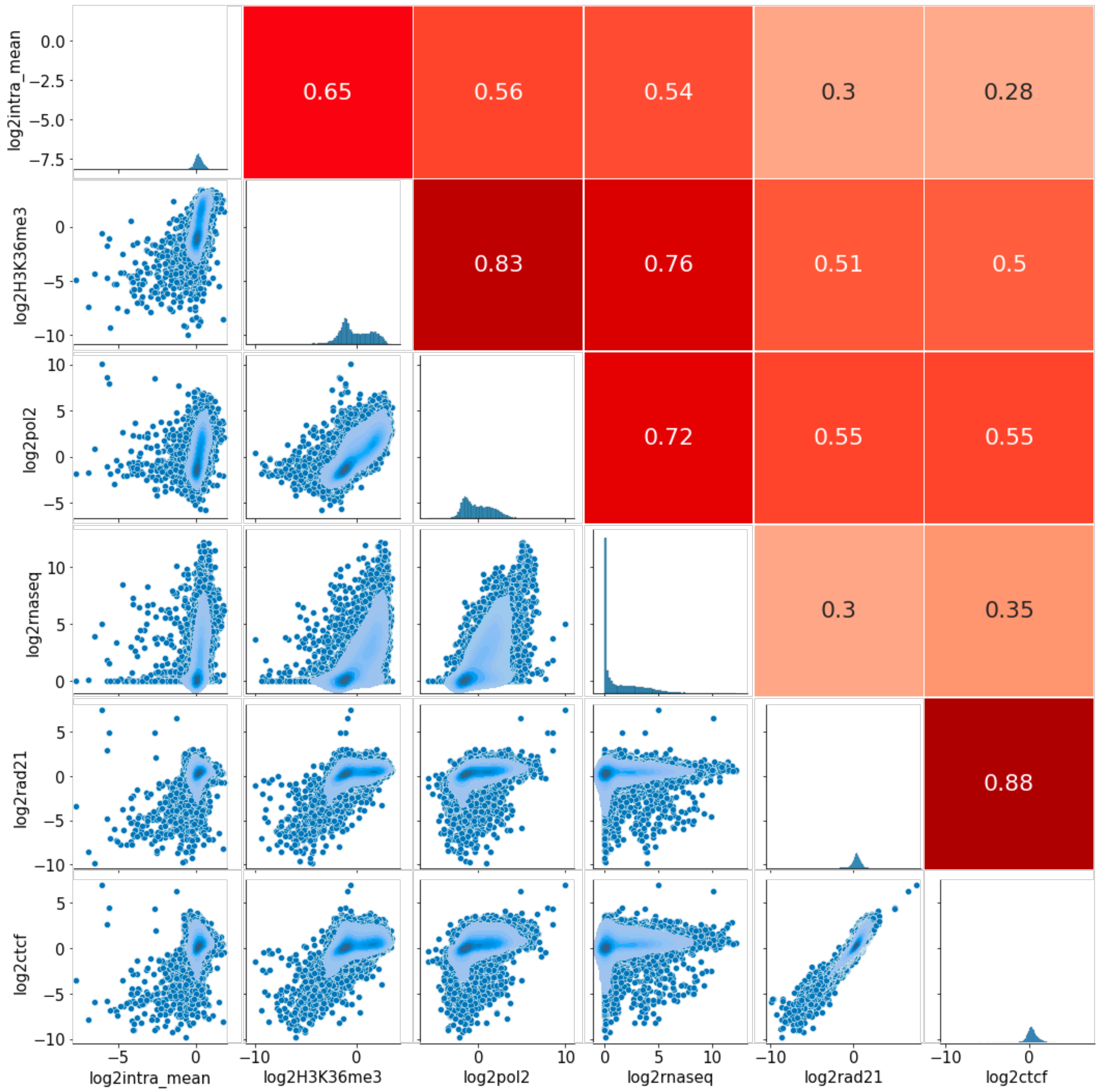


**Fig. S1. Correlations and scatter plots between intragene contact enrichment and various (epi)genomic datasets** (H3K36me3, Pol II, RNA-seq, RAD21 and CTCF).

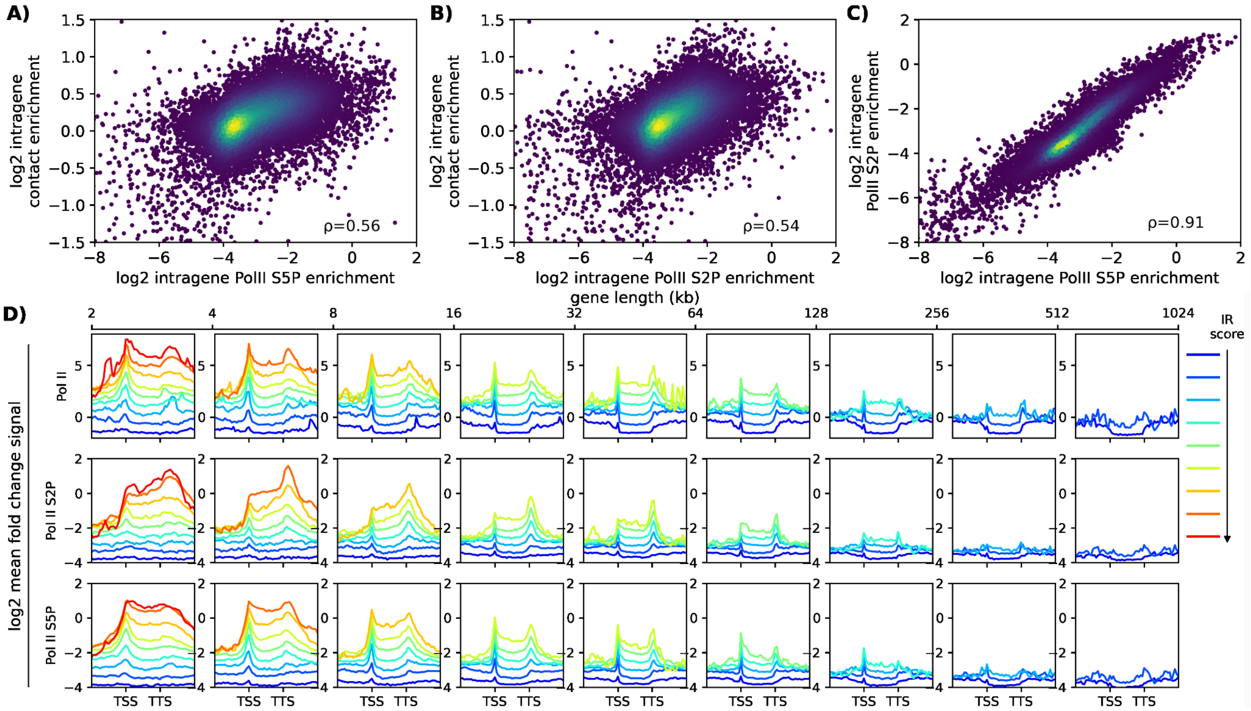


**Fig. S2. Investigating the Impact of RNA PolII Phosphorylation State.** (**A,B**) As in Fig1B of the main text. These scatterplots demonstrate the correlation between intra-gene contact score and the average intra-gene signals for RNA PolII Ser5P (A) and Ser2P (B). (**C**) Scatterplot showing the correlation between average intra-gene signals for RNA PolII Ser5P and Ser2P. Spearman correlations are given. (**D**) Comparison of mean fold change signals for RNA PolII (first row), RNA PolII Ser2P (second row), and RNA PolII Ser5P (third row) within the gene domain and the flanking region. Genes are clustered based on length and IR score; each column represents a different gene size range, with colors indicating IR score ranging from low (blue) to high (red). These results suggest a consistent correlation between IC and IR scores, independent of the phosphorylation state.

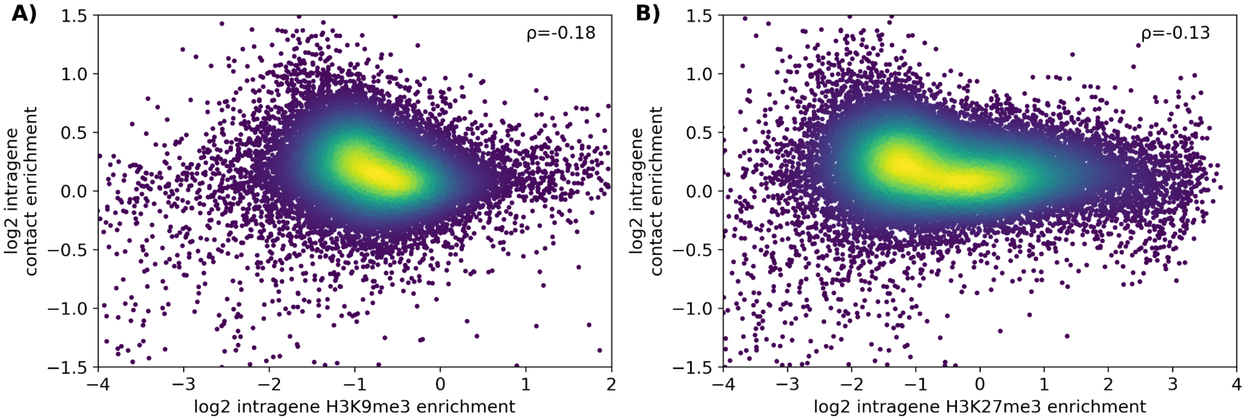


**Fig. S3. Illustration of the negative correlation between intra-gene contact score and repressive histone marks.** As in Fig.1B of the main text. Scatterplots show the correlation between IC score and average intra-gene H3K9me3 (A) and H3K27me3 (B) signals. Spearman correlations are given. The presented data highlight a weak negative correlation between IC score and the repressive histone marks.

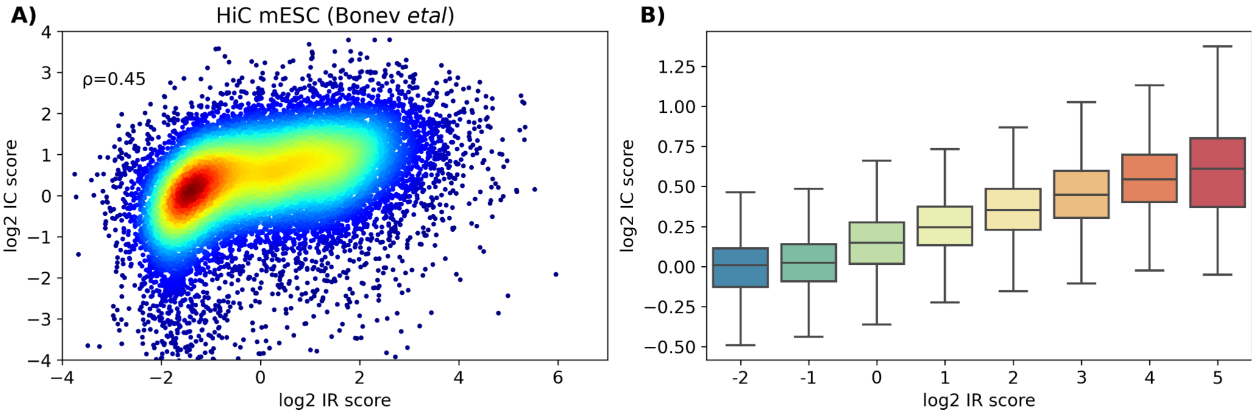


**Fig. S4. Positive correlation between intra-gene contact score and Pol II occupancy from Hi-C experiments on mESC (Bonev et al. 2017).** (**A)** Scatterplot of IC vs IR scores for genes >8kb. Spearman correlation is given. (**B)** Boxplot representation of IC scores after clustering genes with similar IR scores.

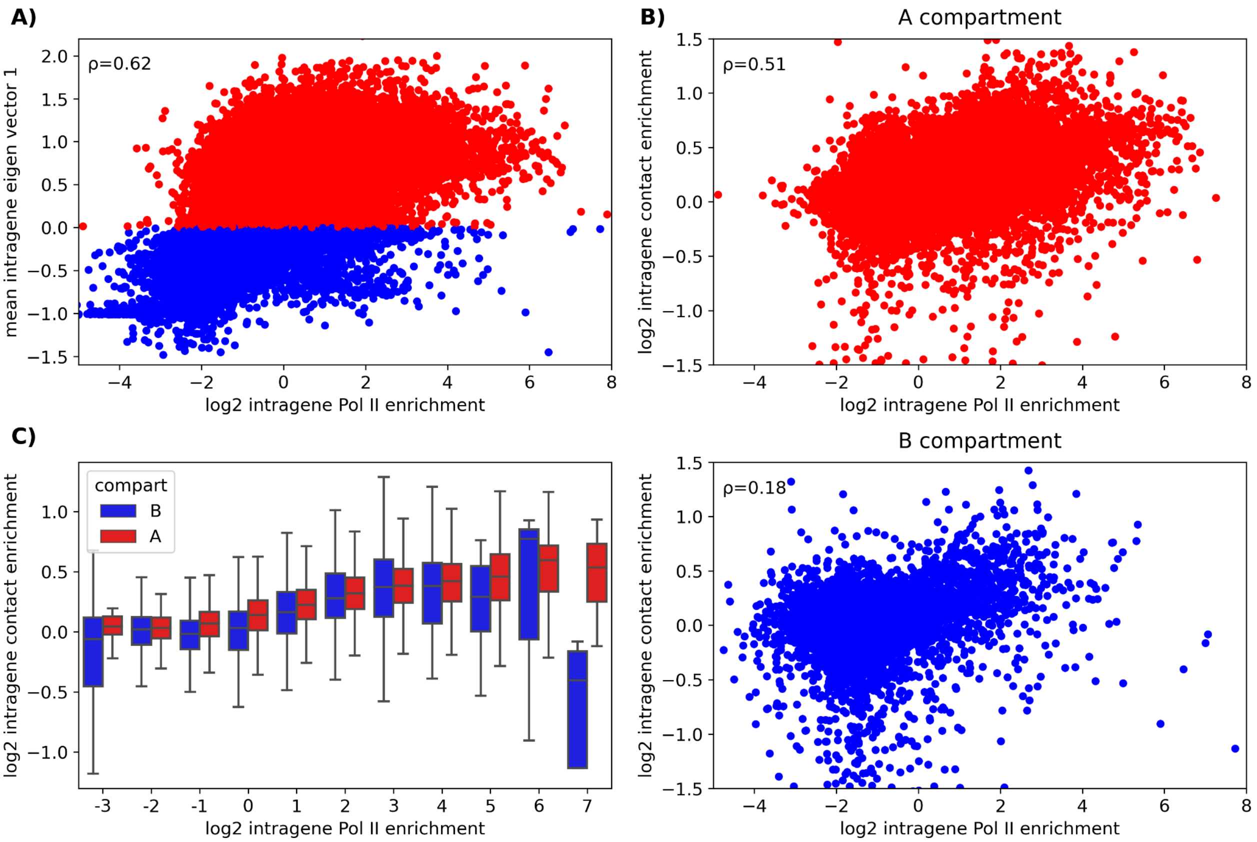


**Fig. S5. Compartment-dependent correlation between intragene contact enrichment and Pol II occupancy. (A)** A positive correlation (spearman's $\rho = 0.62$) between compartment strength (mean value of intragene first eigenvector) and intragene Pol II enrichment (IR). Genes with high (resp. low) Pol II occupancy tend to be in A (resp. B) compartment. (**B)** Correlation and scatter plot between intragene contact enrichment (IC) and IR for genes in A compartment (top) and B compartment (bottom), showing a stronger correlation for genes in A compartment. **(C)** Box plot of IC for different IR values for genes in A or B compartments.

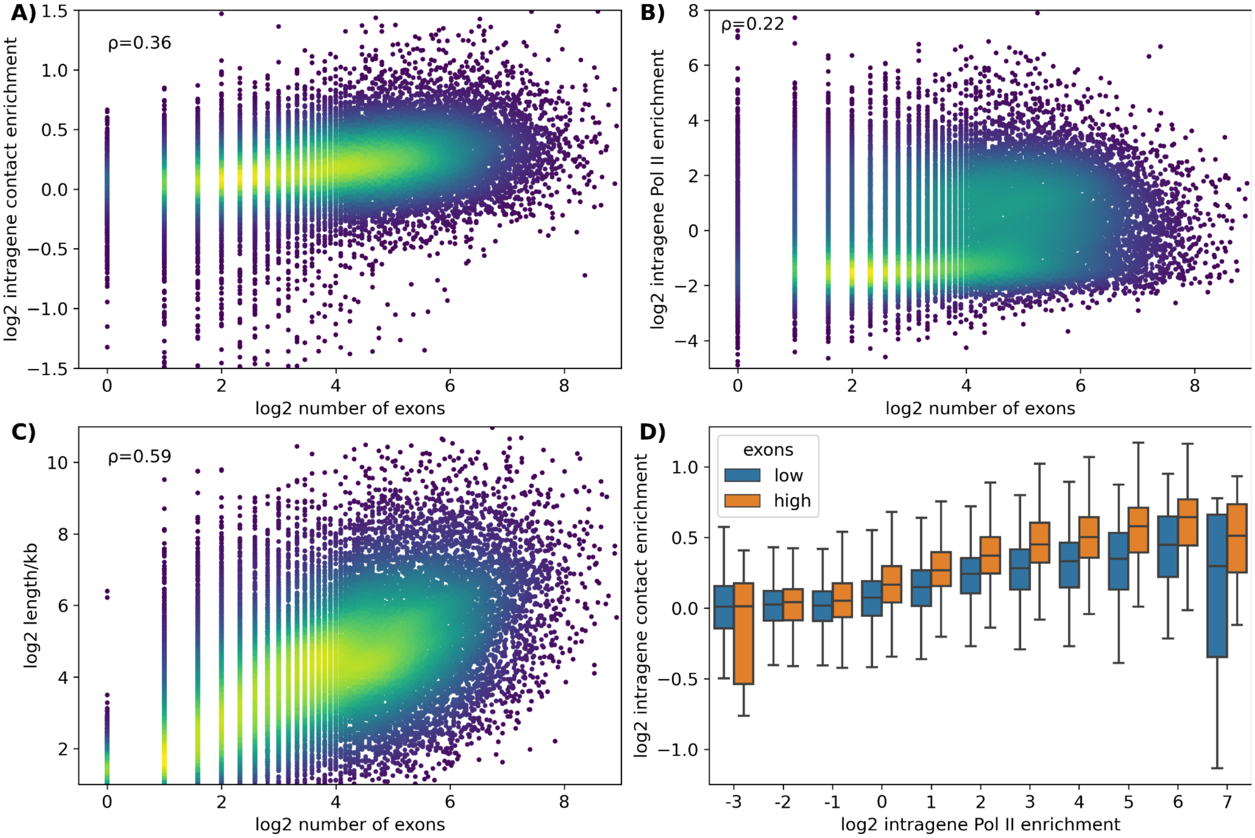


**Fig. S6. RNA splicing may affect gene folding. (A,B,C)** Correlation and scatter plot between number of exons and IC(A), IR (B) and gene length (C). Genes with more exons, and eventually more RNA splicing events, are relatively more compact, occupied by more Pol II and have bigger size. **(D)** Box plot of IC for different IR values for genes with a low or high number of exons (by sorting genes based on the exon numbers). A similar trend is observed for both gene sets, however, genes with a higher number of exons tend to be more compact.

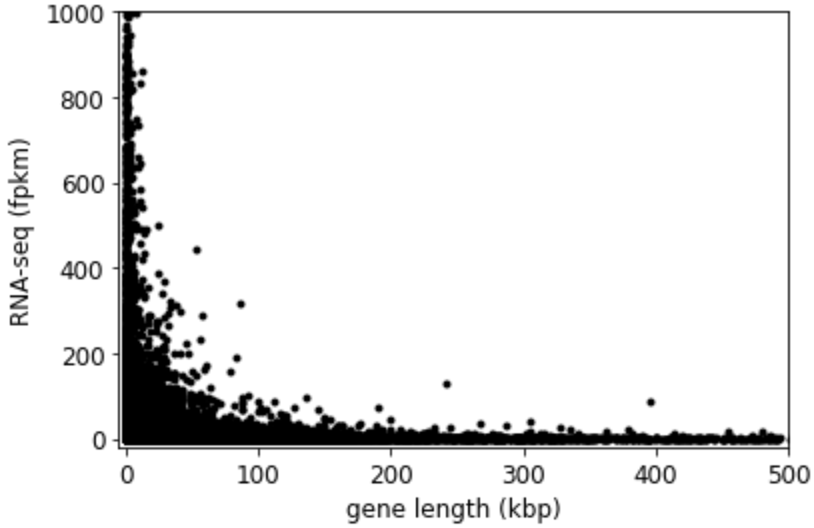


**Fig. S7. Long genes are rarely highly expressed.** Relation between the expression rate (RNA-seq) and gene length for mESC. Most highly expressed genes are short. For instance, there are 1207 short genes (<26kbp) with fpkm>100, compared to only 79 large genes (>26kbp) at the same level of expression.

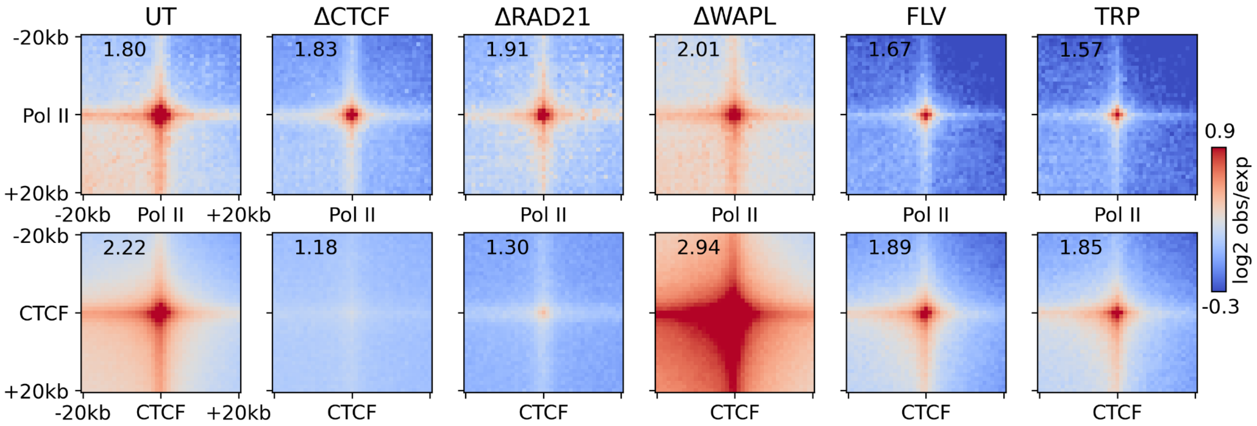


**Fig. S8. Protein-mediated loop structure.** Pile-up analysis of loop enrichment (for pairs of ChIP-seq peaks in distance range of 160-320 kb) around Pol II (top row) and CTCF (bottom row) peaks for wild-type (UT) and different mutants. Enrichments of the pixel intensity at the center relative to the background are given. Pol II-mediated loops vastly remain unchanged after depletion of cohesin loop-extrusion factors (CTCF, RAD21 and WAPL), while for Pol II inhibitors (FLV and TRP) their intensities significantly decrease. On the other hand, CTCF-mediated loops drastically change upon depletion of loop-extrusion factors, while being only slightly perturbed by Pol II inhibitors.

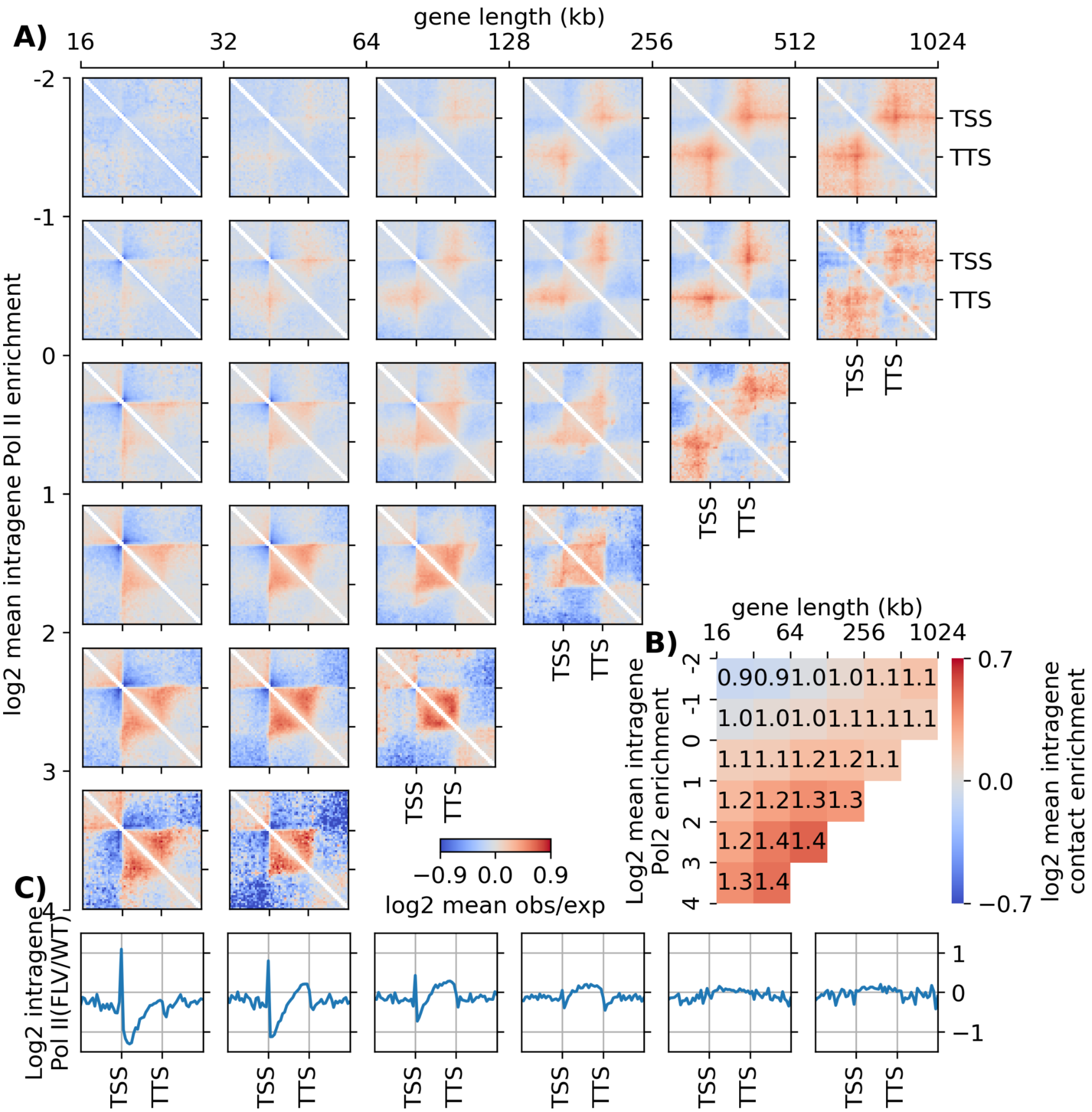


**Fig. S9. Effects of Pol II elongation inhibitor on gene folding**. **(A)** Pileup meta-gene analysis of obs/exp map around genes in FLV-treated cells for clusters based on size (horizontal axis) and Pol II occupancy in WT (vertical axis). Maps for clusters with less than 25 representative genes were not drawn, due to lack of statistics. **(B)** Averaged intragene contact enrichment for each cluster. The color bar is presented in a log2 scale, while the values are represented in a linear scale. **(C)** Averaged occupancy of Pol II along genes in FLV-treated cells relative to wild-type ones. Decreasing the Pol II occupancy in gene body may significantly decrease the intragene folding structure.

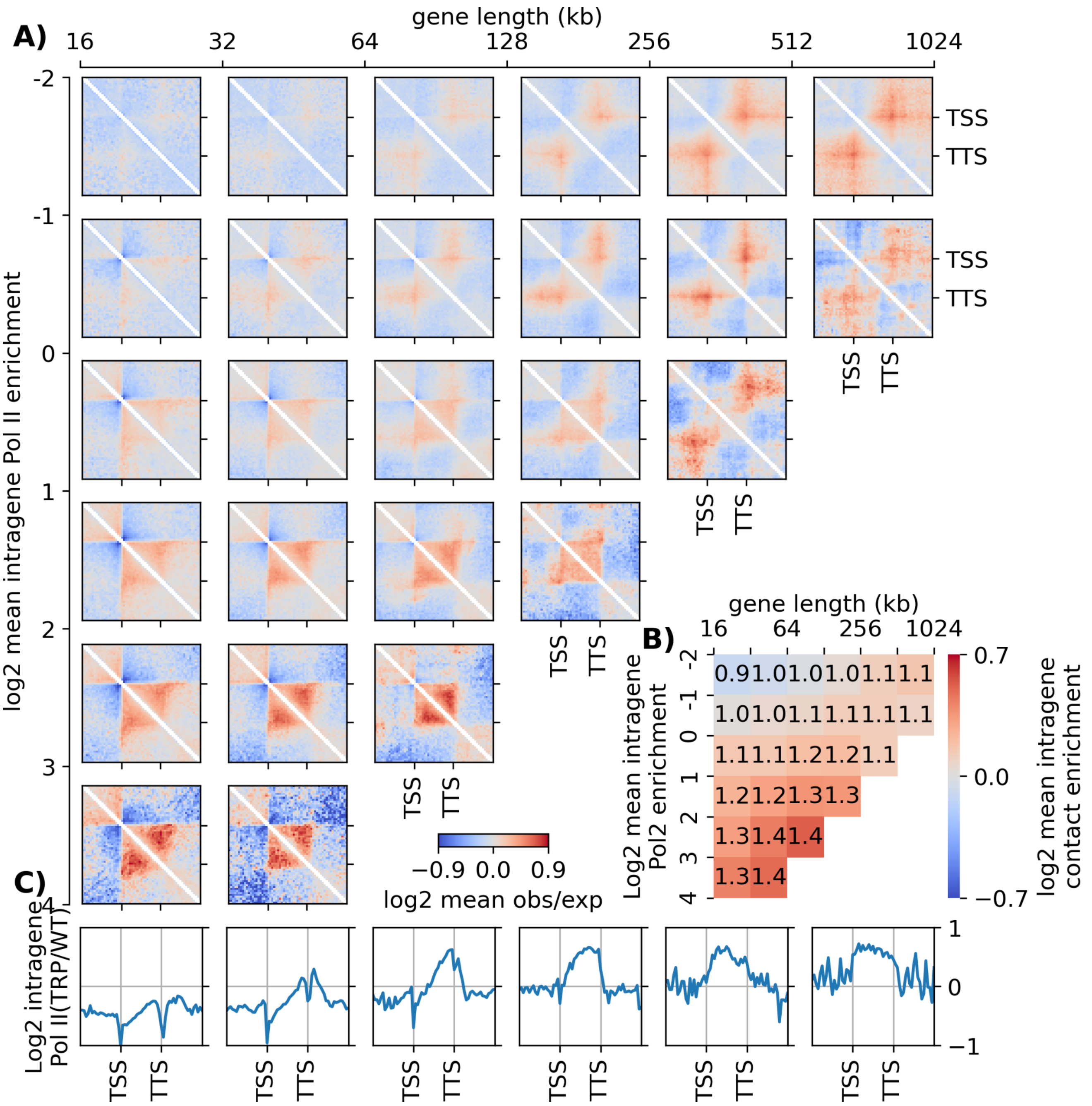


**Fig. S10. Effects of Pol II initiation inhibitor on gene folding**. **(A)** Pileup meta-gene analysis of obs/exp map around genes in TRP-treated cells for clusters based on size (horizontal axis) and Pol II occupancy in WT (vertical axis). Maps for clusters with less than 25 representative genes were not drawn, due to lack of statistics. **(B)** Averaged intragene contact enrichment for each cluster. The color bar is presented in a log2 scale, while the values are represented in a linear scale. **(C)** Averaged occupancy of Pol II in TRP-treated cells relative to wild-type ones. Decreasing the Pol II occupancy in the gene body may significantly decrease the intragene folding structure.

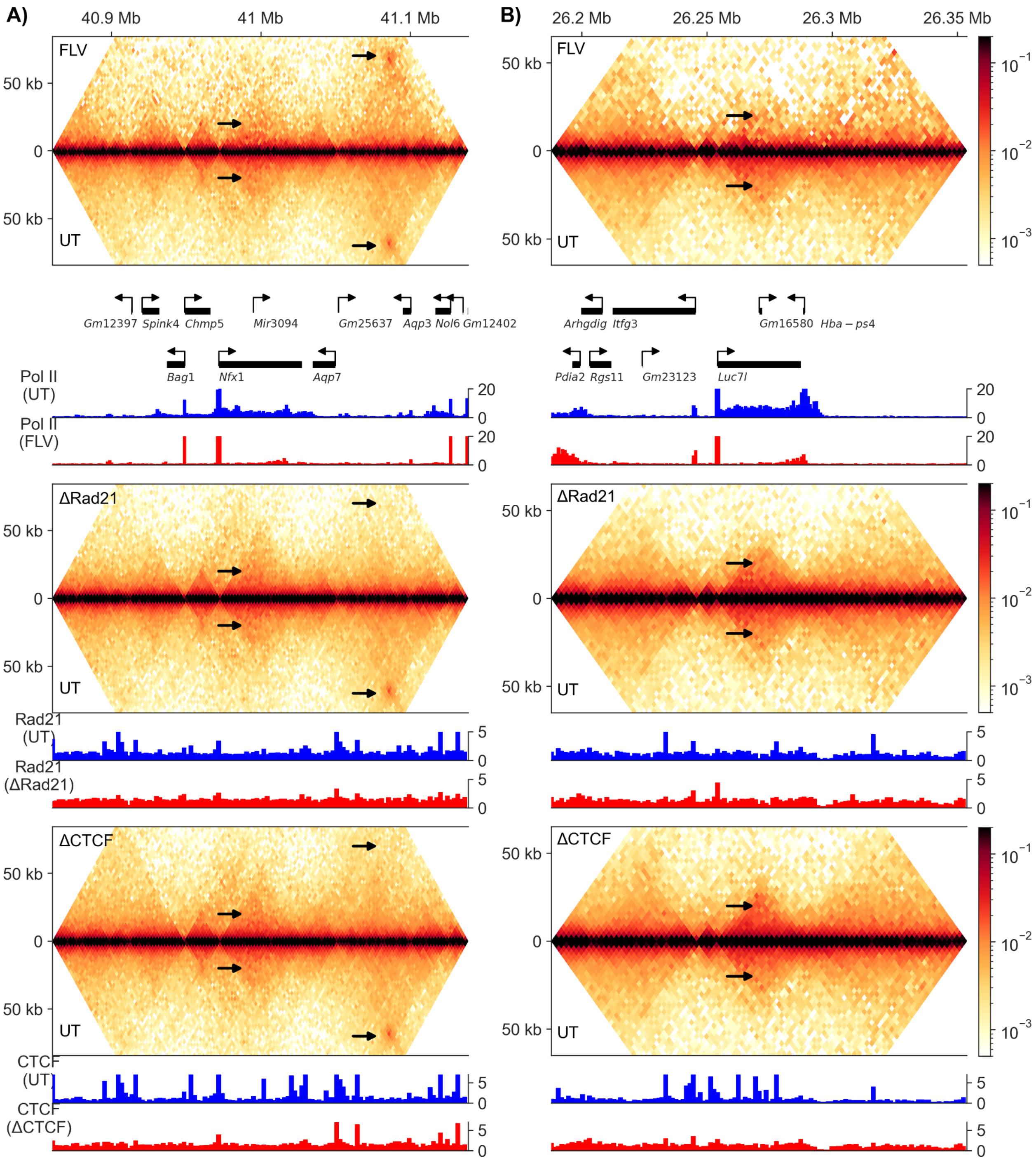


**Fig. S11. Effects of Pol II, cohesin and CTCF on local genome organization. (A,B)** Comparison of contact maps between wild-type and different mutants, FLV (top), RAD21-depletion (middle) and CTCF-depletion (bottom) followed by genes and ChIP-seq tracks for two different regions, (A) chr4:40861-41136 kbp and (B) chr17:26188-26351 kbp. The gene structures are partially disrupted after FLV treatment, while it remains unchanged upon RAD21- and CTCF-depletions. Also, FLV treatment does not alter CTCF-mediated loops, whereas RAD21- and CTCF-depletions remove these loops.

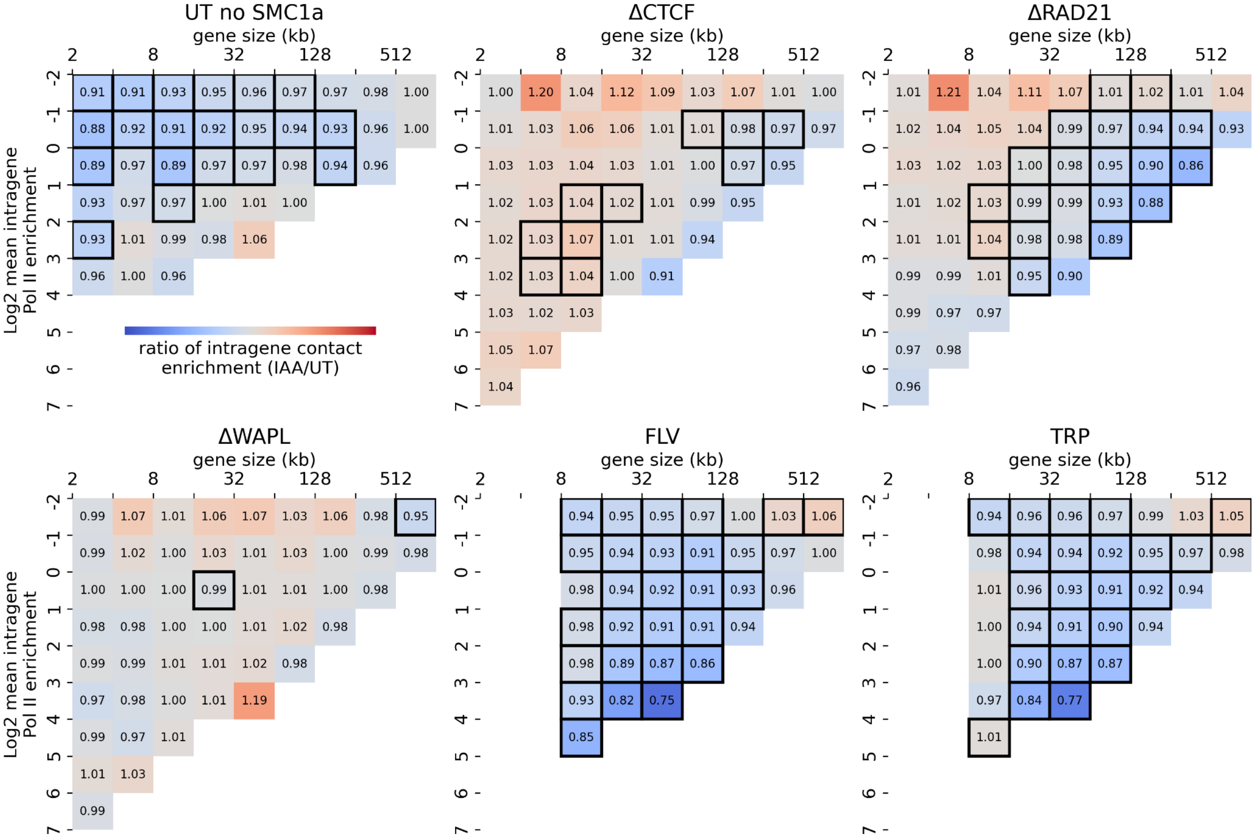


**Fig. S12. Effect of different mutants on intragene contact enrichments.** Heatmaps of the ratio between the averaged intragene contact enrichments of mutant and wild-type for different gene clusters based on gene size (horizontal axis) and Pol II enrichment (vertical axis). The bold squares are indicating the significant changes (*P*-value <0.05). Pol II inhibitors (FLV and TRP) significantly decrease gene folding level, while perturbations in CTCF and cohesin levels have negligible or mild effects on the gene compaction.

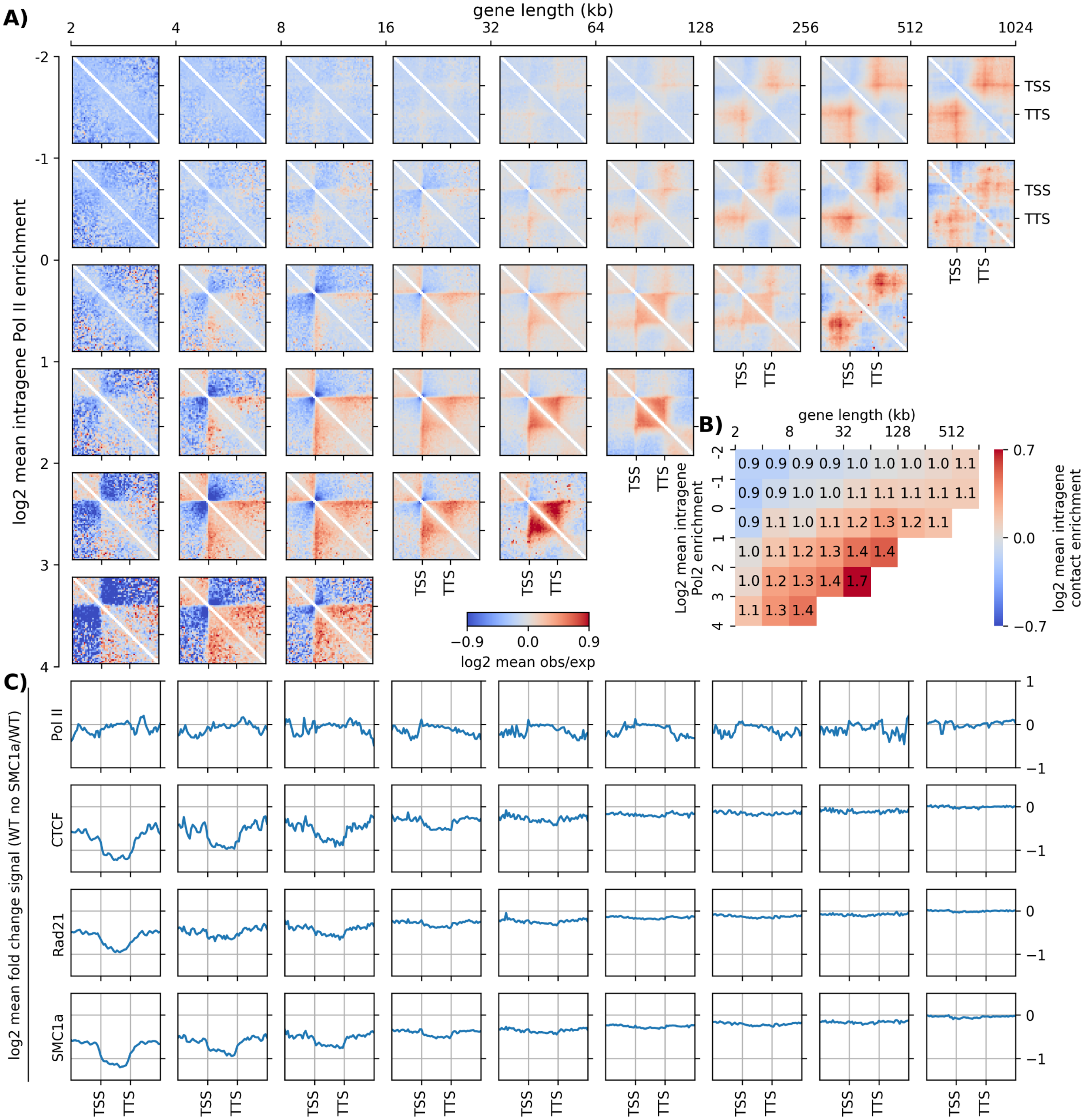


**Fig. S13. Cohesin occupancy of the gene has negligible or mild effect on gene folding**. **(A)** Pileup meta-gene analysis of obs/exp map around genes for clusters based on size (horizontal axis) and Pol II occupancy (vertical axis). These data are for wild-type cells and genes with low cohesin occupancy (genes having a SMC1 occupancy lower than the median value). Maps for clusters with less than 25 representative genes were not drawn, due to lack of statistics. **(B)** Averaged intragene contact enrichment for each cluster. The color bar is presented in a log2 scale, while the values are represented in a linear scale. **(C)** Ratios of averaged ChIP-Seq signals between genes with low cohesin occupancy and all genes for different chromatin binders. Changing cohesin occupancy without changing Pol II occupancy does not change drastically intragene folding structure.

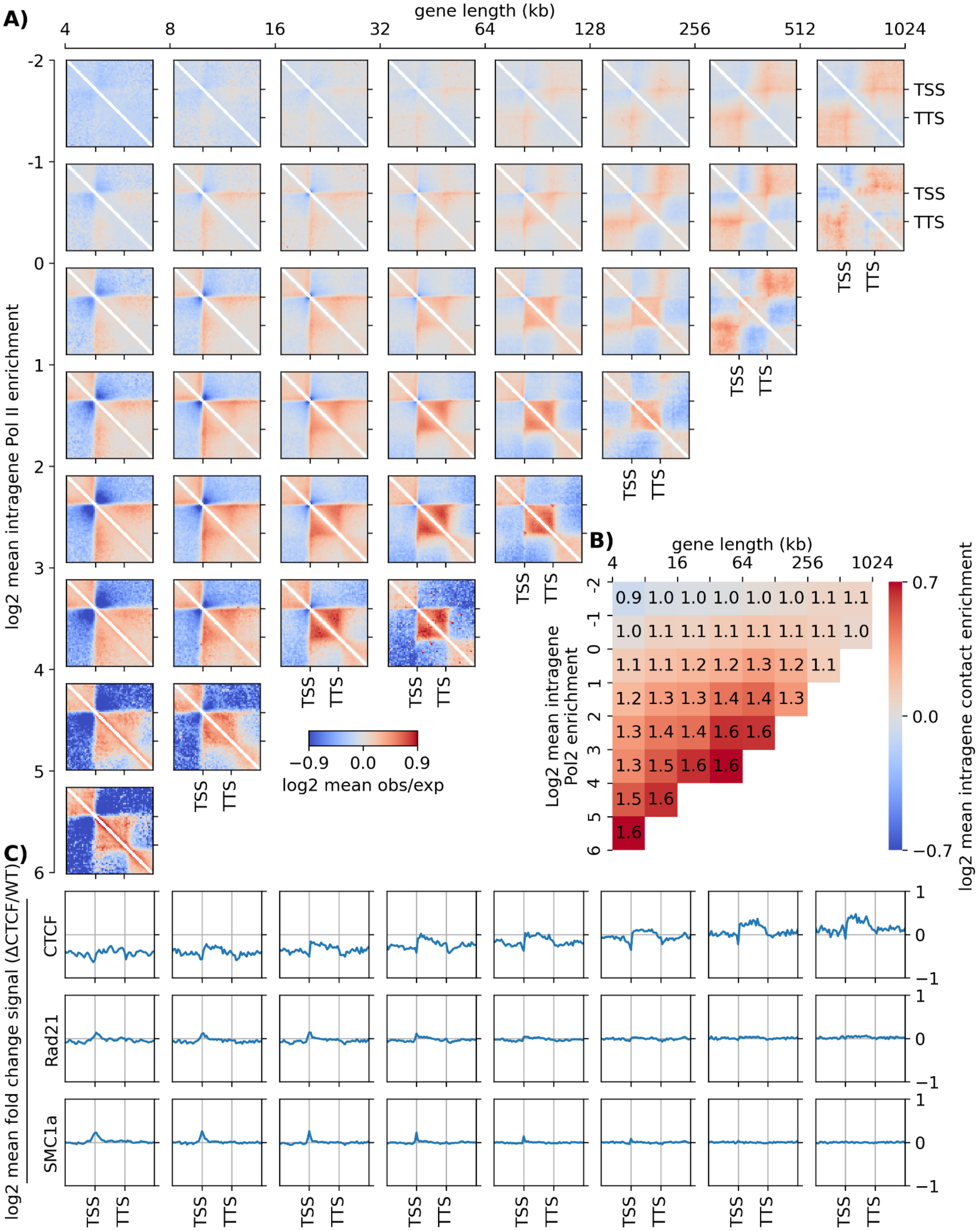


**Fig. S14. Effects of CTCF removal on gene folding**. **(A)** Pileup meta-gene analysis of obs/exp map around genes for clusters based on size (horizontal axis) and Pol II occupancy (vertical axis) for CTCF-depleted cells. Maps for clusters with less than 25 representative genes were not drawn, due to lack of statistics. **(B)** Averaged intragene contact enrichment for each cluster. The color bar is presented in a log2 scale, while the values are represented in a linear scale. **(C)** Averaged occupancy of different chromatin binders for CTCF-depleted cells relative to wild-type ones. Depletion of CTCF may decrease the promoter stripes outside the gene, but it has negligible effect on intragene folding structure.

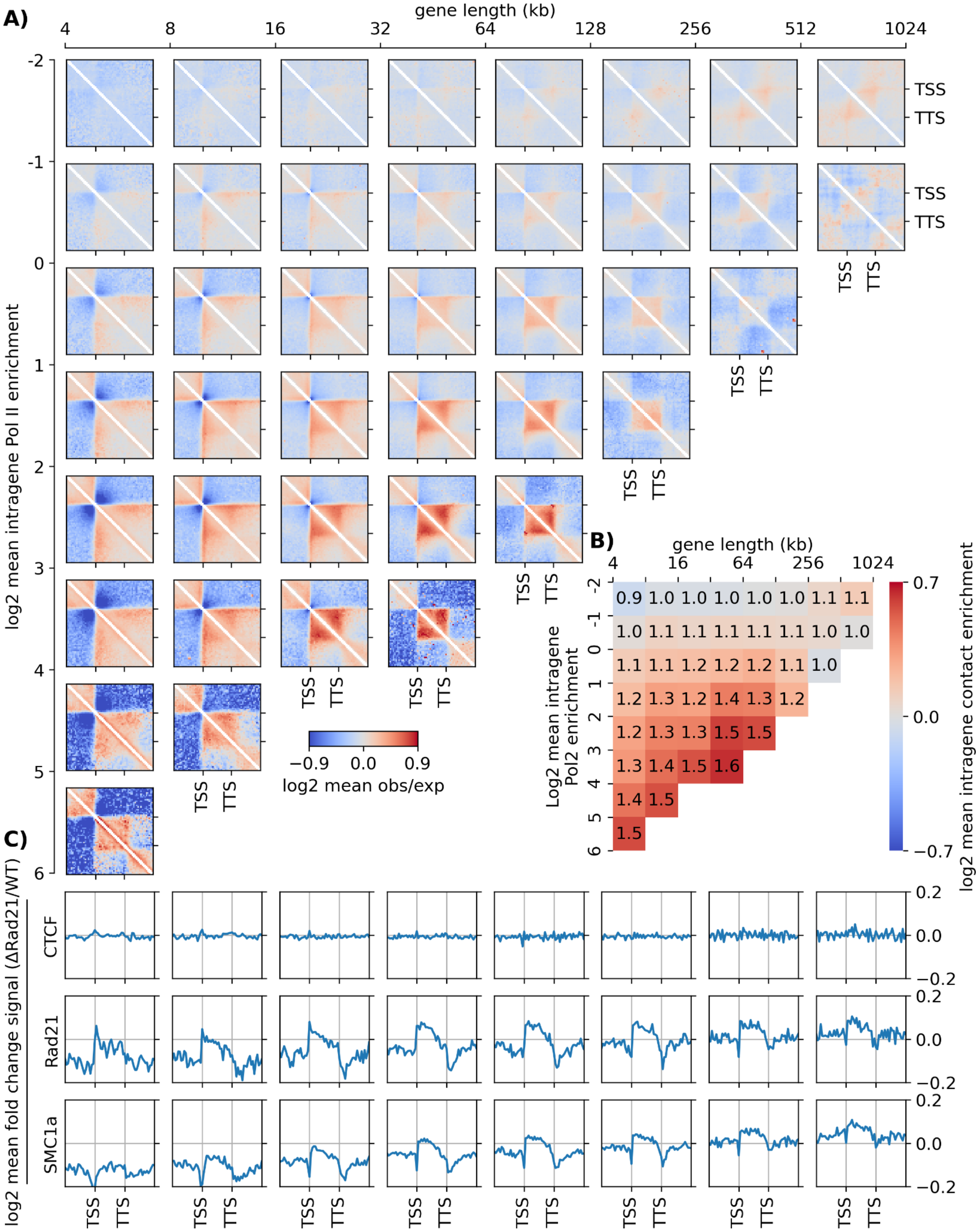


**Fig. S15. Effects of cohesin removal on gene folding**. **(A)** Pileup meta-gene analysis of obs/exp map around genes for clusters based on size (horizontal axis) and Pol II occupancy (vertical axis) for RAD21-depleted cells. Maps for clusters with less than 25 representative genes were not drawn, due to lack of statistics. **(B)** Averaged intragene contact enrichment for each cluster. The color bar is presented in a log2 scale, while the values are represented in a linear scale. **(C)** Averaged occupancy of different chromatin binders for RAD21-depleted cells relative to wild-type ones. Depletion of cohesin may decrease the promoter stripes outside the gene, but it has mild effect on intragene folding structure.

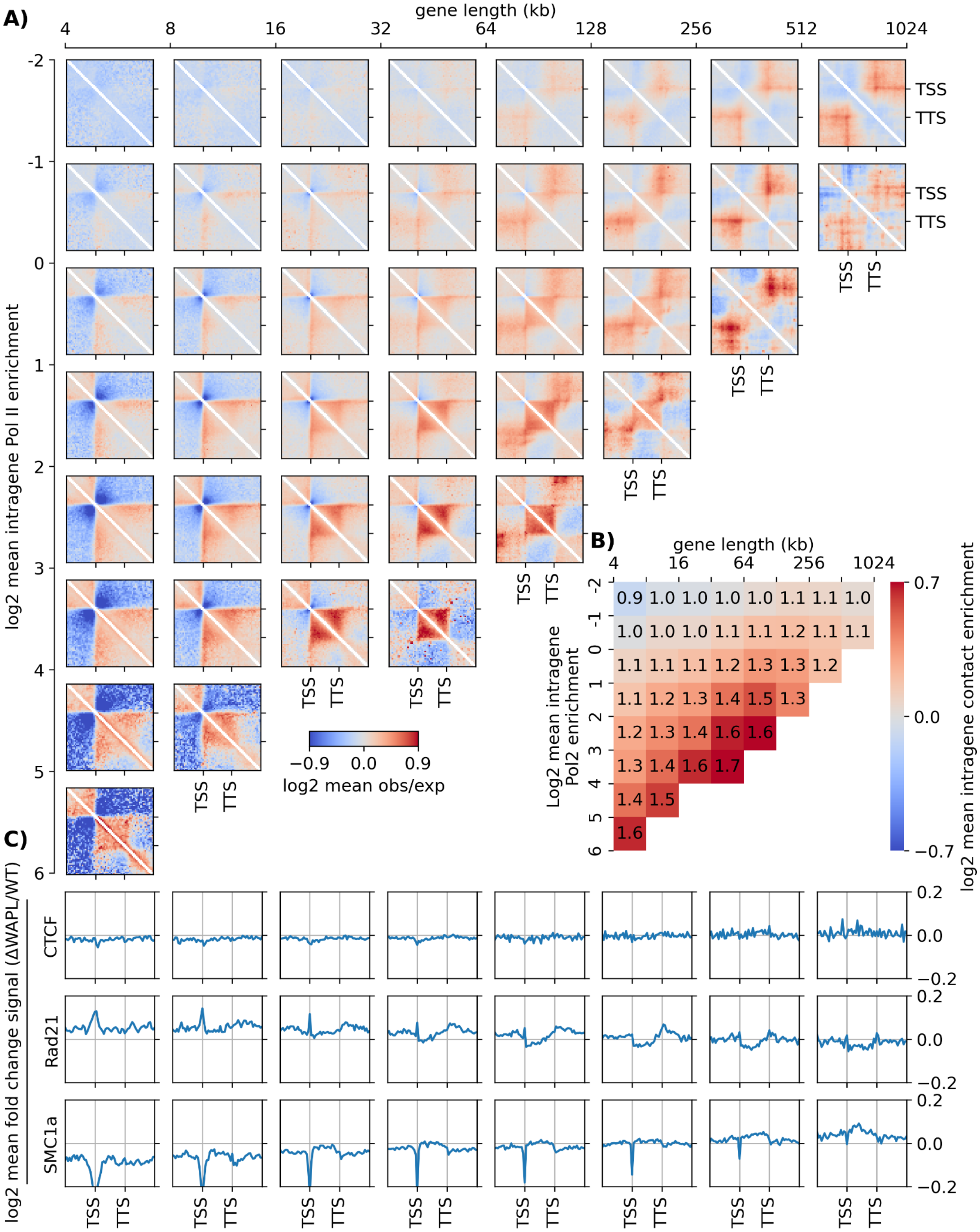


**Fig. S16. Effects of increasing cohesin occupancy on gene folding**. **(A)** Pileup meta-gene analysis of obs/exp map for gene clusters based on size (horizontal axis) and Pol II occupancy (vertical axis) for WAPL-depleted cells. Maps for clusters with less than 25 representative genes were not drawn, due to lack of statistics. **(B)** Averaged intragene contact enrichment for each cluster. The color bar is presented in a log2 scale, while the values are represented in a linear scale. **(C)** Averaged occupancy of different chromatin binders for WAPL-depleted cells relative to wild-type ones. Depletion of WAPL and eventually increasing the cohesin occupancy may increase the promoter stripes outside the gene, but it has negligible effect on intragene folding structure.

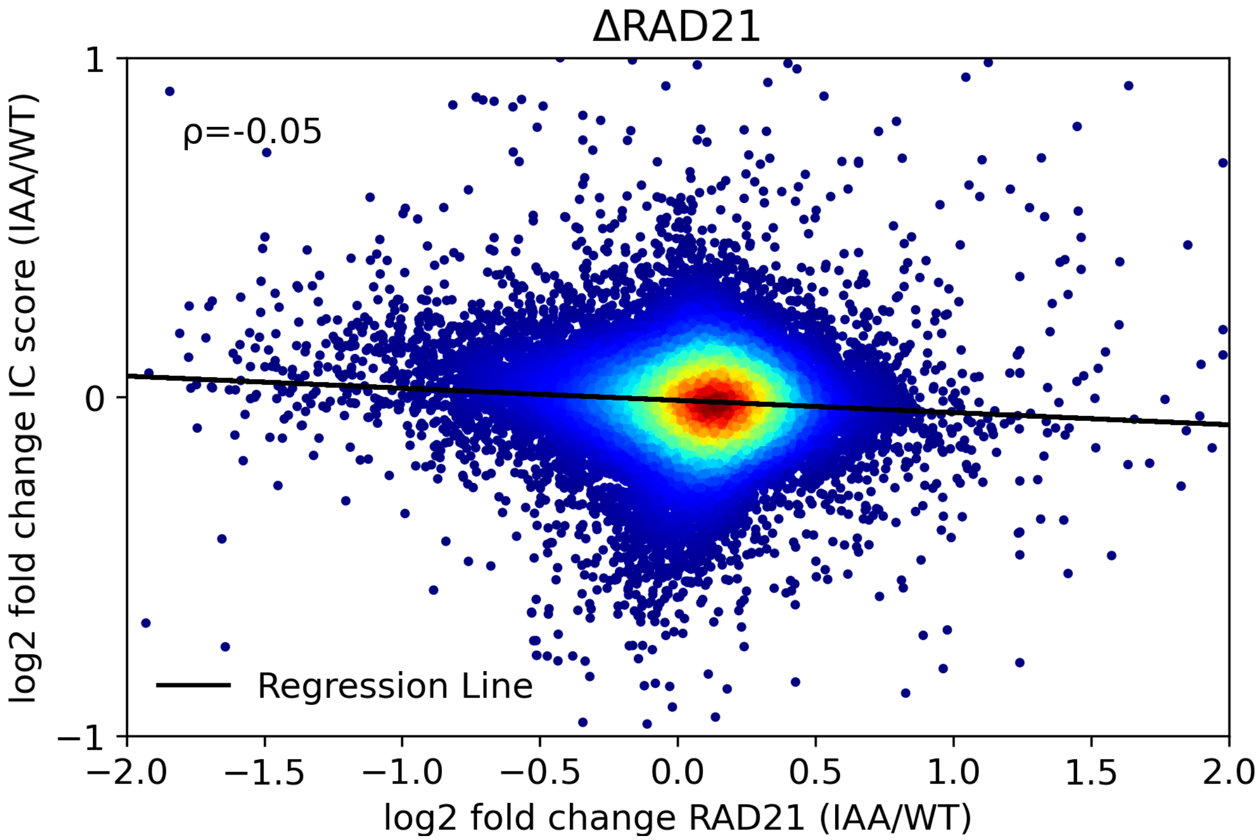


**Fig. S17. Scatter plot of fold change in IC scores vs fold change in IR scores after RAD21 degradation.** No correlation is observed between the change in RAD21 occupancy and changes in IC scores.

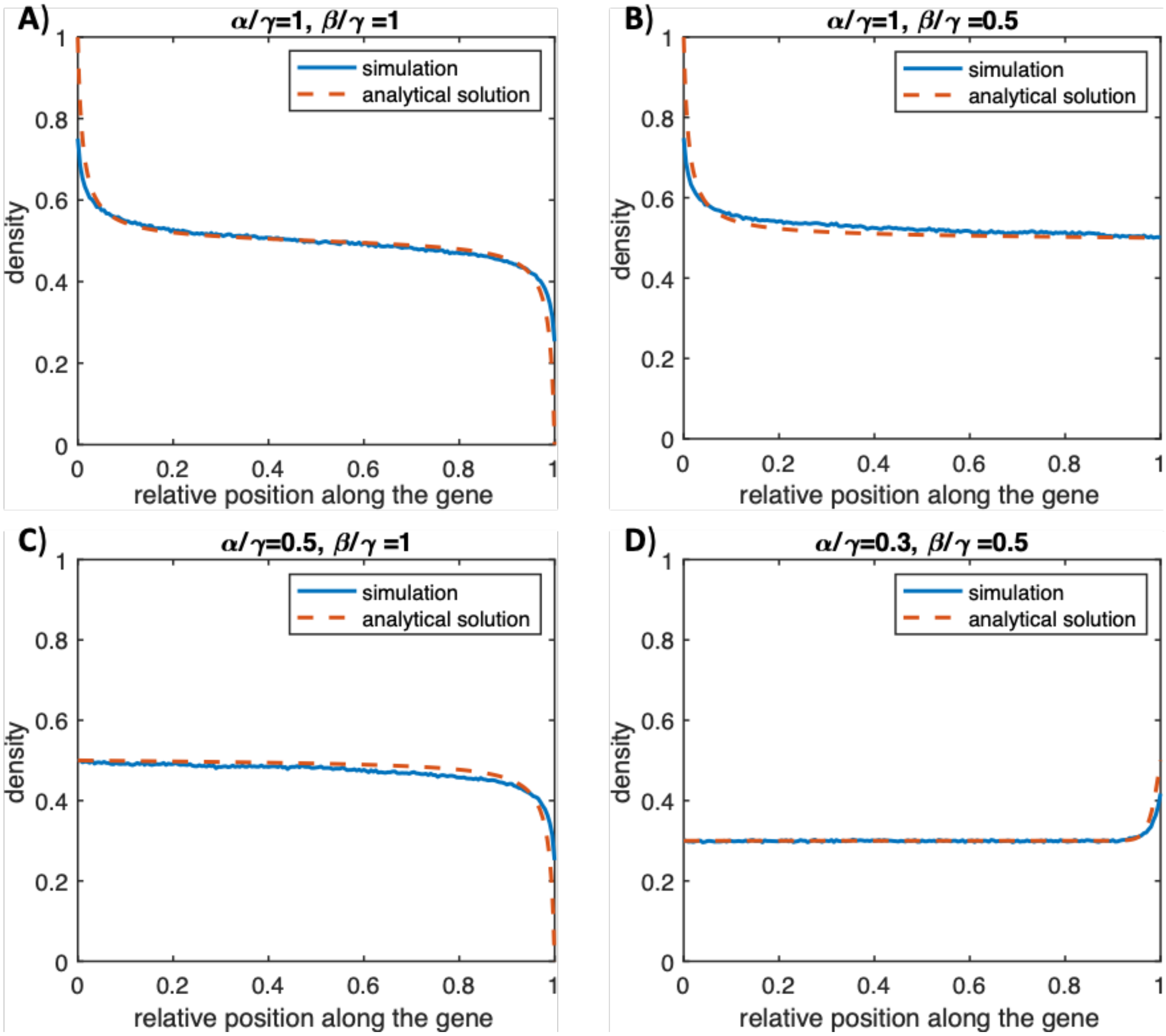


**Fig. S18. Comparison between analytical solution and simulation results of the TASEP model. (A-D)** Predictions of Pol II density along the gene for a simple TASEP model ($\gamma_0 = \gamma$ ) for different parameter sets with a good agreement between analytical solution and Monte-Carlo simulations.

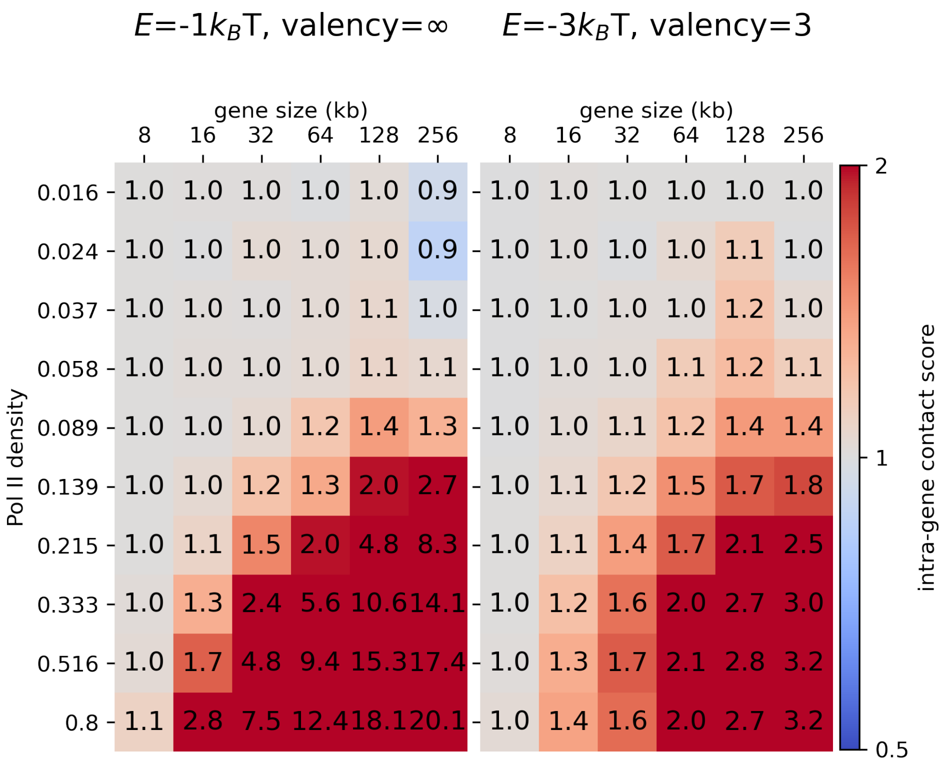


**Fig. S19. Predicted heatmaps of intra-gene contact score against gene size and Pol II density for two different valency numbers and strength of interactions.** We conclude that unrestricted valency even with lower strength of interaction cannot recapitulate quantitatively the behavior observed in experimental data, while for limited valency (2 or 3), model predictions are in good agreement with experiment.

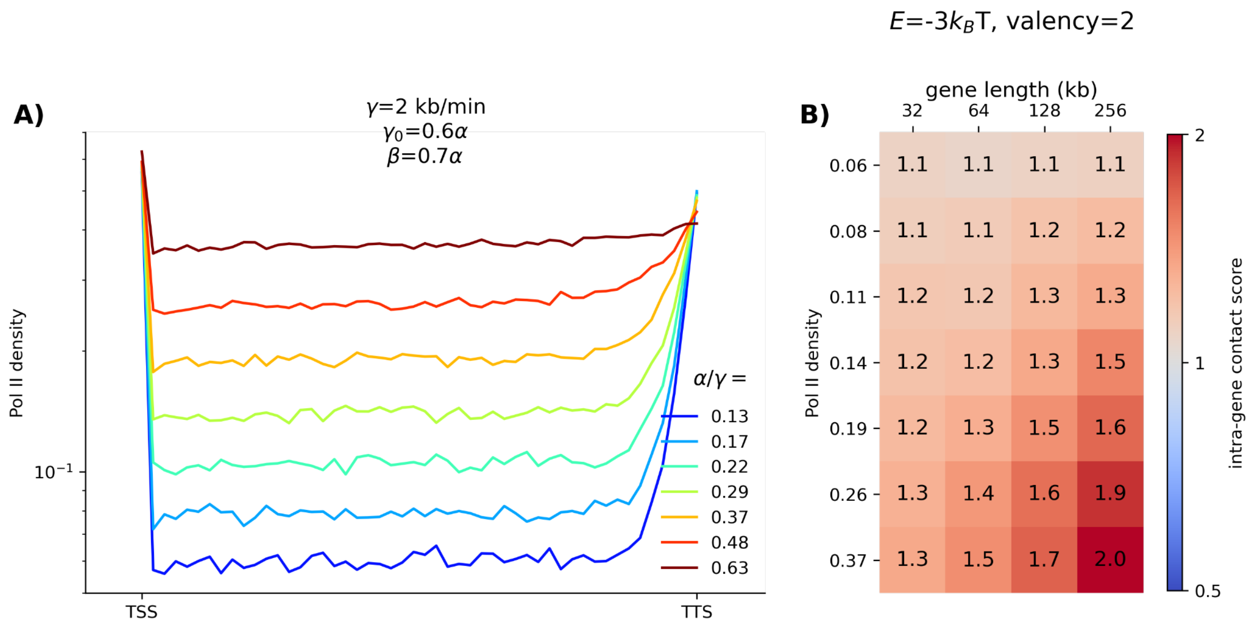


**Fig. S20. Exploring the impact of non-uniform Pol II profiles on intra-gene contact score. (A)** Various non-uniform density profiles of Pol II with significant peaks at the both ends for a fixed elongation rate ($\gamma$=2kb/min) and different binding rates ($\alpha$), with $\gamma_0 = 0.6\alpha$ and $\beta = 0.7\alpha$. (**B)** Predicted heatmap illustrating of intra-gene contact score against gene size and RNA PolII density of panel (A), for E=-3$k_BT$ and valency=2. These data indicate that non-uniform RNA PolII profiles predict a similar correlation between IC score, IR score and gene size when compared to uniform Pol II profiles.

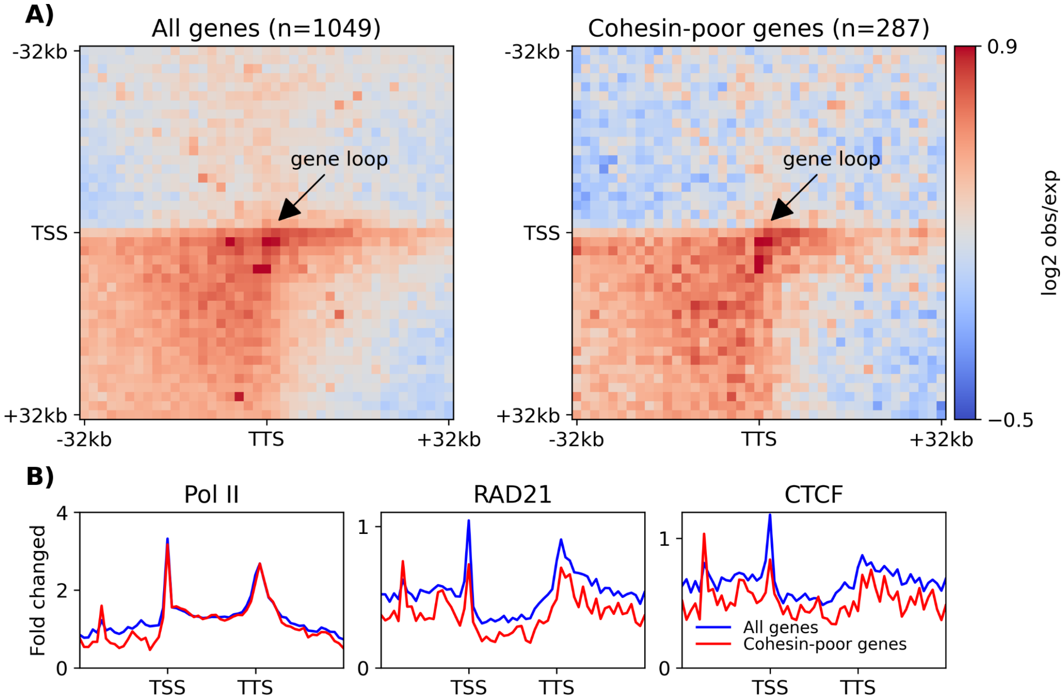


**Fig. S21. Correlation between Pol II profile and gene structure. (A)** TSS-TTS interactions (gene-loop) for long genes (higher than 40kbp) with high Pol II occupancies at TSS and TTS (left) and same for a subset of genes with low RAD21 occupancy (right). (**B)** Comparison of different averaged ChIP-seq profiles in the genes area for the two gene groups mentioned in (A). Gene-loops are less affected by cohesin-mediated loop extrusion and mainly driven by Pol II-Pol II interactions.

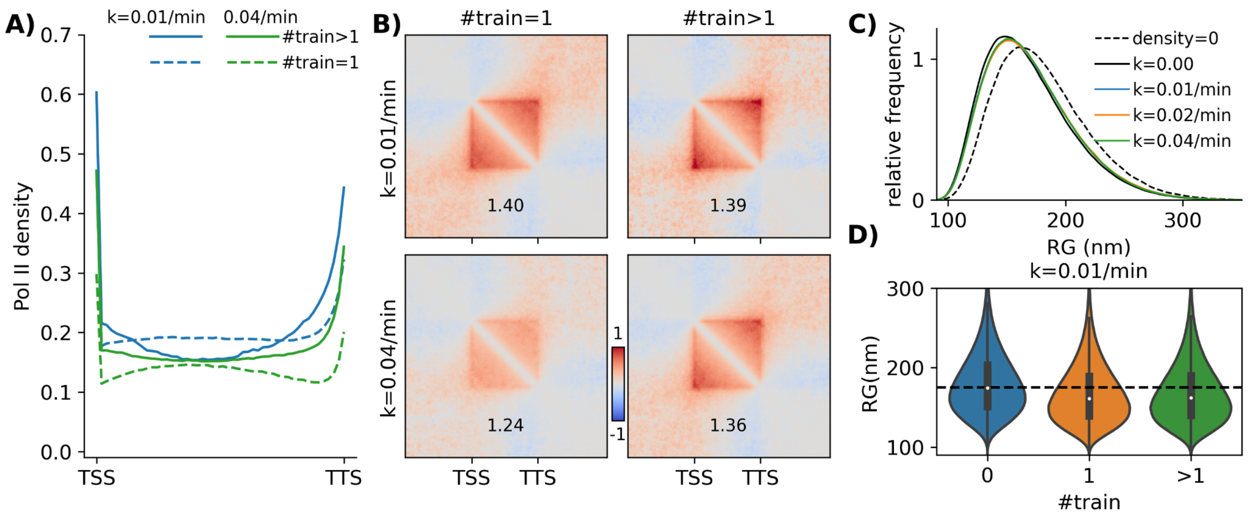


**Fig. S22. RNA Pol II train size may alter gene compaction. (A)** RNA Pol II density profiles when the gene contains one (dashed lines) or more than one (solid lines) trains of Pol II. **(B)** Predicted contact maps for conditions similar to (A). **(C)** Probability distribution functions of the radius of gyration for the null case (no Pol II, black dashed line) and four other different cases with different burst frequencies. **(D)** Violin Plots of the radius of gyration for different train numbers on the gene for k=0.01/min. The black dashed line corresponds to the case without Pol II.

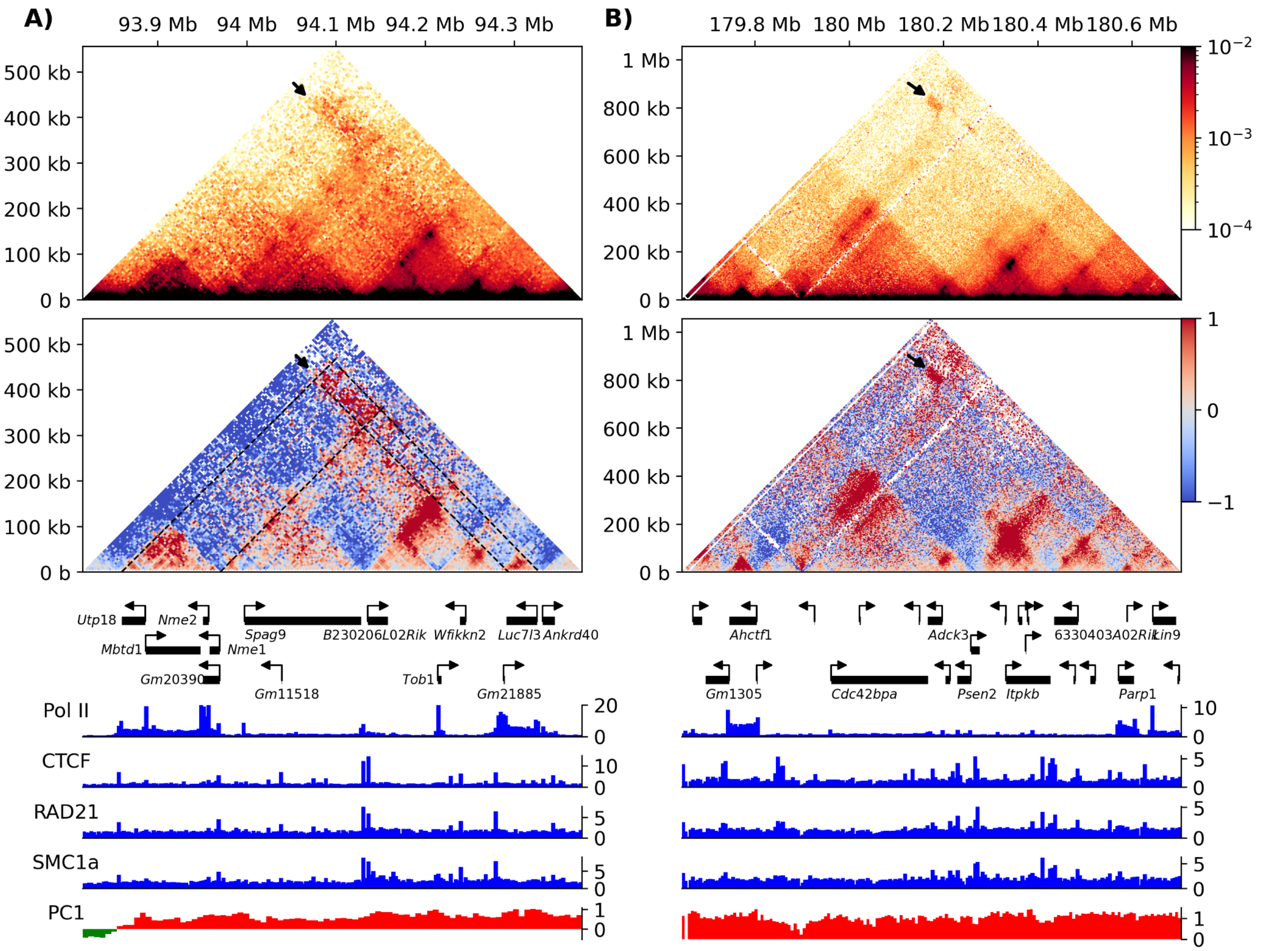


**Fig. S23. Intergene contact between active genes. (A,B)** Contact maps (top) as well as obs/exp maps (bottom) around two different regions, (A) chr11:93816-94372 kbp and (B) chr1:179645-180701 kbp. Gene positions and different ChIP-seq and compartment tracks are given below. Arrows show the significant intergene interactions. Distal domains with high Pol II occupancy may interact together and form a transcriptionally active sub-compartment.

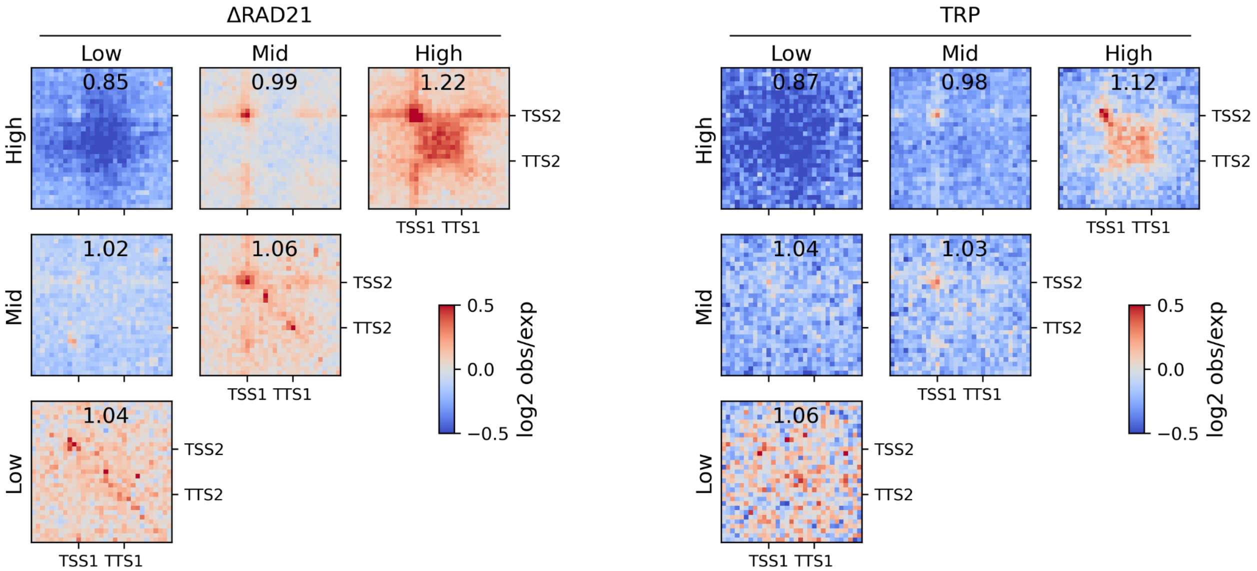


**Fig. S24. PGMA of inter-gene interactions for 32-64 Kb long genes, which are separated by genomic distances between 128kb and 2 Mb, for both RAD21-depleted (left) and TRP-treated (right) cells.** Genes were clustered into three distinct populations (Low, Mid and High) based on their IR scores. The ratios of average inter-gene contact enrichment to background signal are given.

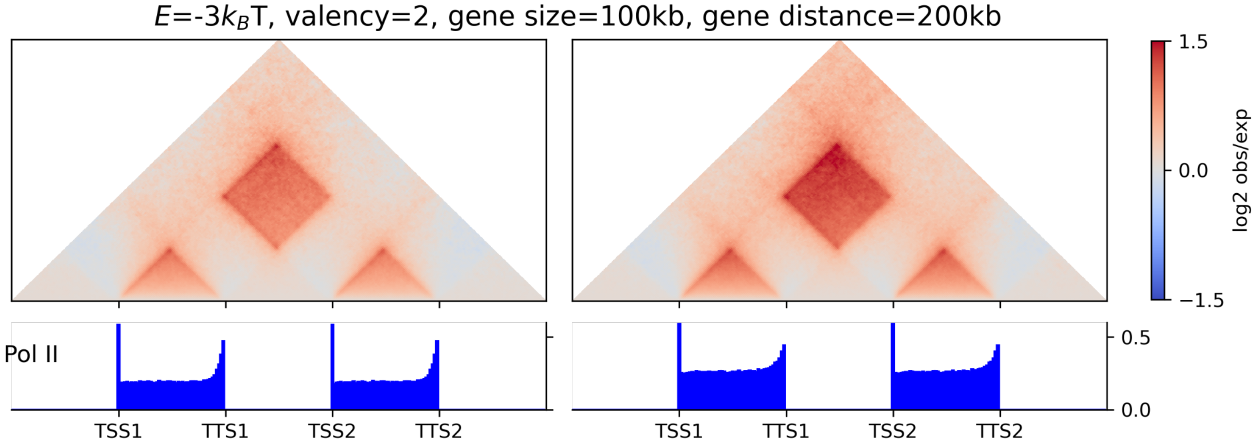


**Fig. S25. Modeling predictions for inter-gene interactions. (Top)** Predicted obs/exp maps of inter-gene interactions between two genes, each with a size of 100kb, and a distance (middle to middle) 200kb, under condition of E=-3$k_B T$ and valency=2. (**Bottom)** Pol II profiles corresponding to top panels.

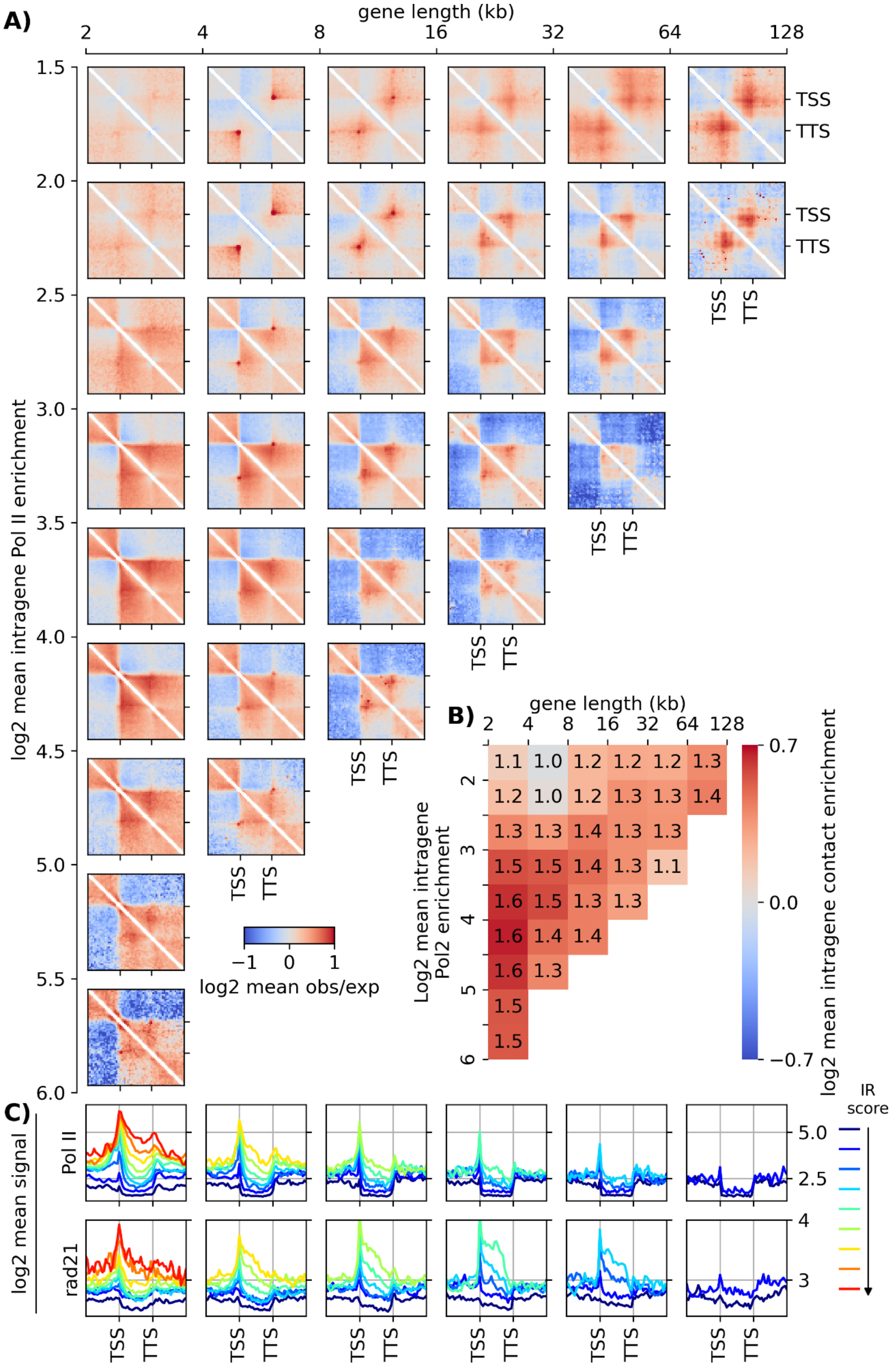


**Fig. S26. Gene classification of *Drosophila*.** **(A)** Pileup meta-gene analysis of obs/exp map around genes for clusters based on size (horizontal axis) and Pol II occupancy (vertical axis) for *Drosophila* nc14 cells. Maps for clusters with less than 25 representative genes were not drawn, due to lack of statistics. **(B)** Averaged intragene contact enrichment for each cluster. The color bar is presented in a log2 scale, while the values are represented in a linear scale. **(C)** Averaged Pol II and RAD21 occupancies for different gene clusters (blue to red are indicating low to high intragene Pol II occupancies).

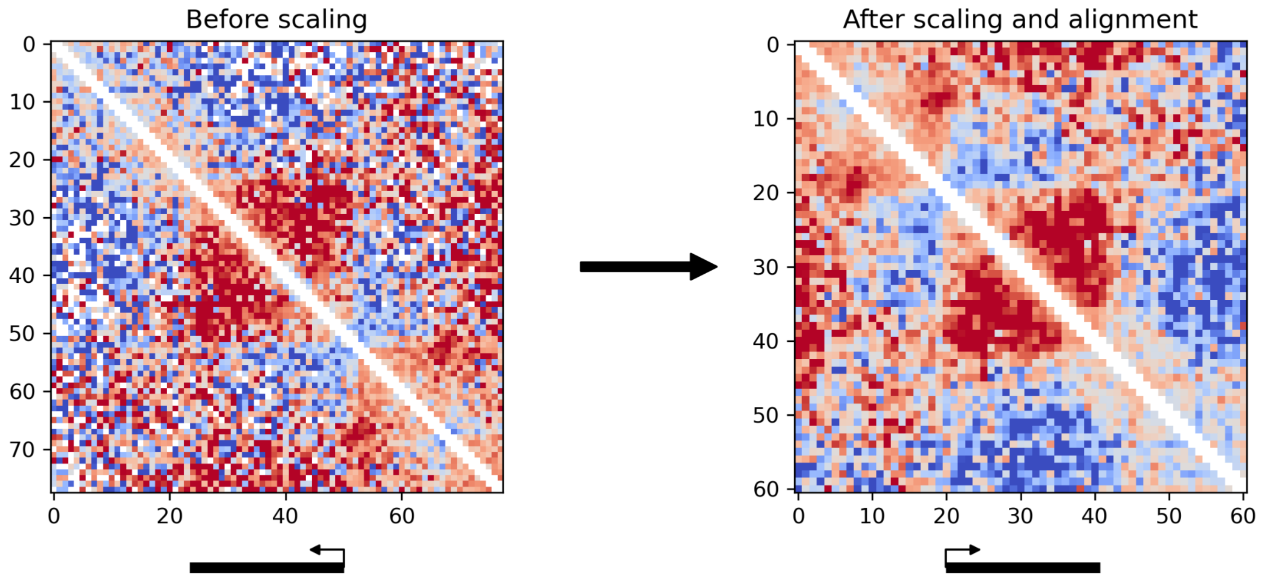


**Fig. S27. An example of rescaling and alignment in pileup meta-gene analysis.** The obs/exp maps of a region chr1:54943-55070kbp, which contains the 42kb-long *Sf3b1* gene in its center, before and after scaling. The resolution before scaling is 1.6kbp, which implies 26 pixels for the gene (and 78 pixels total region). The 78×78 matrix was rescaled to a 60×60 pseudo-matrix with gene meta-size of 20 pixels, and also flipped to align the gene in the forward direction. All the structural details are well conserved after rescaling.

### Supplementary movies

**Movie S1**. Movie showing a typical simulated trajectory of a 100-kb long gene (white monomers) along with two 100-kb long flanking regions (dark blue monomers). The red monomers represent RNA Pol II-bound loci within the genes. The average Pol II density is 0.5, E=-3kT and no restriction on interaction valency. The left panel shows the 3D organization of all the monomers while the right panel only shows the arrangement of Pol II-bound monomers. The lower panel provides a 1D representation of the location of Pol II-bound monomers within the gene. The movie depicts the condensation of Pol II-bound monomers, which subsequently facilitates the compaction of the full gene.

**Movie S2**. As in Movie S1 but for a valency number 2. The movie illustrates how a smaller valency number destabilizes Pol II-mediated condensation, leading to a reduction in intra-gene compaction.

**Movie S3**. Movie demonstrating how bursty gene activity impacts on the structural dynamics of the gene. In the left panel, a typical simulated trajectory for a 200-kb long gene (Cyan monomers) with flanking regions (dark blue monomers). Monomers of different colors represent Pol II-bound regions belonging to different transcriptional trains. The right panel visualizes the spatial arrangement of Pol II-bound monomers from different trains, providing insights into their interplay. The lower panel shows the 1D arrangement of Pol II-bound monomers within the gene, highlighting their positioning. This movie illustrates how gene conformation is influenced and shaped by the dynamic intra- and inter-train interactions.

**Movie S4**. Movie illustrating inter-gene interactions. In the left panel, a simulated trajectory showing the behavior of two distinct genes (white and light blue monomers) with their surrounding genomic regions (dark blue monomers). Pol II-bound loci for each gene are represented by red and orange monomers respectively. The left panel provides insight into the spatial arrangement of Pol II-bound monomers belonging to different genes. The lower panel shows the 1D representation of the location of Pol II-bound monomers within these genes. This movie illustrates how intra- and inter-gene interactions are driven by dynamic Pol II-mediated interactions, ultimately leading to the formation of transcriptionally-active subcompartments.

### Supplementary Notes

#### 1. Analytical solution of TASEP model

Here, we present an analytical solution for TASEP model in a simple case where initiation rate is equal to elongation rate (i.e. $\gamma_0 = \gamma$ ). We assume that the state of monomer $i$ ($s_i$) is 1 if it has a Pol II and 0 otherwise. Therefore, one can write the equation for time-evolution of ensemble averaged of $< s_i >$ as following,

$$d < s_i >/dt = \gamma < s_{i-1}(1 - s_i) > -\gamma < s_i(1 - s_{i+1}) >$$

where $s_{i-1}$ and $s_{i+1}$ are the state of monomers $i - 1$ and $i + 1$, respectively. If we assume that the state of each monomer is independent on the state of their neighbors then the equation can be rewritten as,

$$d < s_i >/dt = \gamma < s_{i-1} >< (1 - s_i) > -\gamma < s_i >< (1 - s_{i+1}) >.$$

In the continuum limit, one can write $< s_i >\equiv \rho(x,t)$ and make the Taylor development:$< s_{i\pm1} > \equiv \rho(x,t) \pm n^{-1}\partial_x\rho(x,t) + (2n^2)^{-1}\partial_x{}^2\rho(x,t) + \mathcal{O}(n^{-3})$, where $\partial_x$ is partial derivative against position coordinate $x$. Therefore, the master equation in the continuum limit would be as,

$$\partial_t\rho(x,t) \approx \gamma n^{-1}\{(2n)^{-1}\partial_x{}^2\rho(x,t) + (2\rho(x,t) - 1)\partial_x\rho(x,t)\},$$

where, the higher order of $\mathcal{O}(n^{-3})$ have been ignored. In the steady state ($\partial_t\rho(x,t) = 0$), we end up with the differential equation:

$$(2n)^{-1}\partial_x{}^2\rho(x) + (2\rho(x) - 1)\partial_x\rho(x) = 0,$$

with two boundary conditions of $\rho(0) = \alpha/\gamma$ and $\rho(n) = 1 - \beta/\gamma$, which refers to binding and unbinding rates at the two ends of the gene. It can be shown that, for the condition of $\alpha/\gamma = 1 - \beta/\gamma$, the differential equation implies a uniform density of $\rho = \alpha/\gamma$ along the gene. We can also numerically solve the differential equation for other conditions and compare it to the results of Monte-Carlo simulations (**Fig S16**).

#### 2. Effective dependency of the strength of Pol II-Pol II interactions

In the main text, we discussed that limiting the valency of interactions to low numbers allows simultaneously to recapitulate quantitatively both the intensity and non-monotonic shape of the correlation between intra-gene compaction (IC) and Pol II enrichment (IR). An alternative and/or complementary, more trivial, possibility could be that the strength *E* of interactions between Pol II-bound monomers is directly dependent on the local or global density of Pol II along the gene. Indeed, assuming that the Pol II self-attractions in the model are effective and translate interactions between bound Pol II and the same third party (e.g., splicing condensates, Mediator or nuclear speckles) (Guo et al, Nature 2019), several mechanisms may lead to an effective dependency of *E* to Pol II occupancy: (1) at high concentration, RNA was recently shown to destabilize transcription-related condensates (Henninger et al, Cell 2021), therefore, the presence of many transcribing Pol II interacting with such a third party may increase the local concentration of RNAs and, beyond a given threshold, may undermine the stability of the third-party condensates ; (2) the number of sites in third-party condensates accessible to Pol II may be limited, thus beyond a given density of Pol II, competition to interaction sites may occur between transcribing Pol IIs, leading to a lower probability to be bound to the condensate and thus to effectively interact with other Pol IIs.

In both examples, *E* would remain constant as a function of Pol II density until a given threshold above which it would be decreasing for larger Pol II occupancy. If the decrease is strong enough to compensate for the increase of Pol II-bound monomers, this would also lead to a non-monotonic shape for the IC vs IR curve.
